# Airway immune signatures of protection and disease progression in recent human tuberculosis household contacts

**DOI:** 10.64898/2026.01.29.702551

**Authors:** William J. Branchett, Jee-Whang Kim, Jessica Shields, Probir Chakravarty, Jo Lee, Ismail Novsarka, Hubert Slawinski, Katalin A. Wilkinson, Robert J. Wilkinson, Anver Kamil, Raman Verma, Pranabashis Haldar, Anne O’Garra

**Author notes:** Equal first/last authors (William J. Branchett, Jee-Whang Kim, Pranabashis Haldar, Anne O’Garra). Correspondence to: Dr Anne O’Garra.

## Abstract

Whilst the majority of individuals infected with *M. tuberculosis* control the infection and remain asymptomatic, only 5-10% progress to active tuberculosis (TB)^1^. Previous or current infection with *M. tuberculosis* is detected using antigen-specific interferon (IFN)-γ release assays (IGRA), which cannot identify those who will remain healthy or progress to active TB^1^. However, ^18^F-Fluorodeoxyglucose positron emission-computed tomography (PET-CT) can detect increased immune cell metabolic activity in lung parenchyma and intrathoracic lymph nodes associated with infection^2^. The local early immune factors dictating protection or disease progression have not been defined. To address this, we interrogated the airway immune response at single cell resolution in bronchoalveolar lavage (BAL) from extensively clinically characterised recent household contacts of TB patients, who either controlled the infection or progressed to TB disease. Using unbiased analysis of bulk and scRNA-seq of BAL samples, we define type I IFN-dependent and -independent neutrophil signatures in active TB patients and contacts that progressed to TB. We additionally report an inverse relationship between airway neutrophils and T cells, with T cells showing signatures of exhaustion, cytotoxicity and cell death in progressors and TB patients with a neutrophil dominated airway profile. Conversely, T cell signatures of protection in contacts who remained healthy were dominated by genes related to regulation, quiescence and a stem-like profile. We show that both the inflammatory neutrophil signature of TB progression and the stem-like T cell signature of non-progressors from human airways were recapitulated in scRNA-seq data from non-human primate (NHP) granulomas, associated with disease or immune protection, respectively. Our findings from early human airway responses in TB contacts reveal genes, pathways and cell states that may dictate infection outcome and inform strategies for host-directed therapy and vaccine studies.

## Main

A quarter of the global population is estimated to have been infected with *Mycobacterium tuberculosis*, the causative pathogen of tuberculosis (TB). However, only 5-10% of infected individuals progress to active TB^1^, typically within 1-3 years^3,4^. The early local immune factors that determine protection or disease progression are unclear. Bacillus Calmette-Guérin (BCG), the only licensed TB vaccine, offers poor protection against pulmonary TB in adults, highlighting the need to better understand mechanisms of immune protection against human TB^5-7^. Past or current infection with *M. tuberculosis* is detected based on peripheral memory T cell responses using an antigen-specific interferon (IFN)-γ release assay (IGRA), however this does not distinguish current from past infection or predict progression to active TB. Detection of subclinical lung pathology by ^18^F-Fluorodeoxyglucose positron emission and computed tomography (PET-CT), which reveals highly metabolically active cells, has better positive predictive value for clinical progression, consistent with evolving pulmonary and intrathoracic nodal inflammatory changes during progression^2,8^. Blood transcriptomics have identified neutrophil driven type I interferon (IFN) inducible signatures in active TB that correlate with disease severity and progression, supporting a role for this pathway in failed immunity and immunopathology of TB^4,9,10^. The role of type I IFN in exacerbated TB disease is supported by studies in experimental models^11-15^. Control of *M. tuberculosis* is dependent on T cell-mediated immunity^16^, but the precise mechanisms underpinning this are unclear^6,7^.

Key questions remain regarding: (1) the mechanisms of failed immune control of *M. tuberculosis*, which could inform strategies for host-directed immunomodulating therapies; and (2) the early, local, immune signatures associated with control of *M. tuberculosis* infection, which could provide correlates of protection and strategies for improving TB vaccines. Here, we performed in-depth studies of the lower airway immune response in recent household contacts of TB patients in a low TB burden setting where the *M. tuberculosis* re-exposure risk is low, unlike in high TB burden countries^17^, to reveal the primary immune events accompanying protection or disease progression.

### Characterising the airway immune response in TB patients and contacts

To identify the early airway events which may dictate whether recent household contacts of pulmonary TB patients progress to TB or remain healthy, bulk and single cell RNA-sequencing (RNA-seq), flow cytometry and multiplex protein assays were performed on bronchoalveolar lavage (BAL) samples from 11 contacts who progressed to TB and 55 who did not progress, termed non-progressors (23 IGRA^+^ and 32 persistently IGRA^-^) (**Fig. 1a; Extended Data Fig. 1a-b; Supplementary Table 1a,c**). Progression of 11 contacts, out of a total of 153 contacts, was consistent with expected rates^18^ and was accompanied by a positive thoracic PET-CT signal that increased with progression (**Extended Data Fig. 1c, Supplementary Table 1a, Supplementary Fig. 1a**). A stable positive PET-CT signal was detected in a subset of IGRA^+^ but not IGRA^-^ non-progressor contacts (**Extended Data Fig. 1c, Supplementary Table 1a, Supplementary Fig. 1b-c**). All BAL data were compared to those from active pulmonary TB patients at diagnosis (**Fig. 1a; Extended Data Fig. 1a; Supplementary Table 1b**). Unbiased analysis of BAL bulk RNA-seq distinguished TB patients and progressing contacts from non-progressors and was not skewed by age, sex or ethnicity (**Extended Data Fig. 2a-b and Supplementary Table 2**). Expression of genes related to inflammation, T cell and neutrophil functions was increased in BAL from progressors and active TB patients (**Extended Data Fig. 2b and Supplementary Table 3).**

**Fig. 1.**
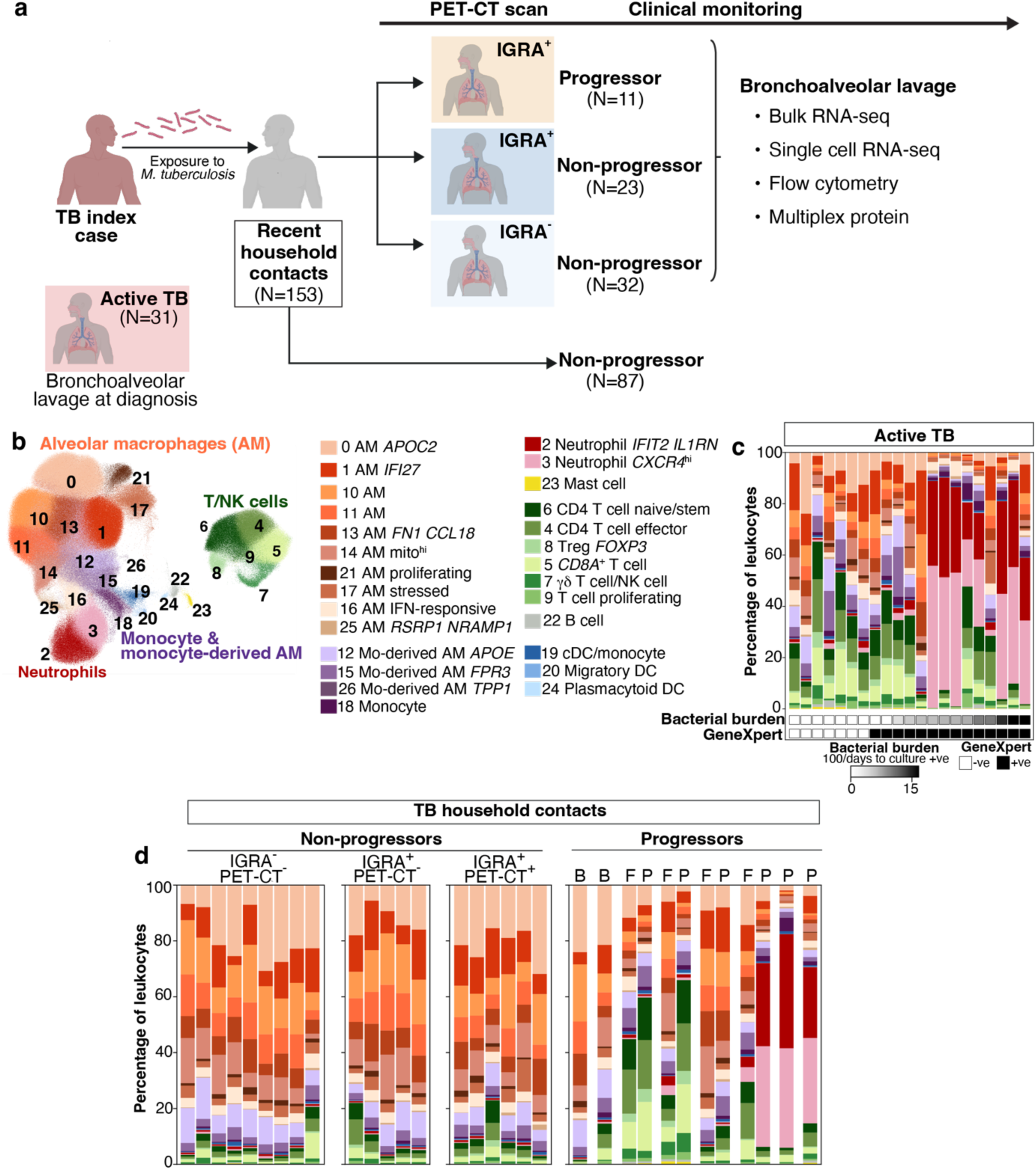
The local airway immune response in TB patients and household contacts. **a.** Scheme of the TB household contacts study. Further details are available in the **Methods, Extended Data Fig. 1a-b** and **Supplementary Tables 1a-c**. **b.** UMAP visualization of total leukocytes (894,159 cells) from all TB patient and household contact BAL samples (total 77 samples from 62 individuals). **c-d.** Stacked bar plots showing percentages of cells in each scRNA-seq cluster per participant. **c.** Active TB patients (N=21) are ordered by increasing BAL *M. tuberculosis* bacterial burden (100/days to culture positivity, with a value of 0 indicating negative BAL culture), with *M. tuberculosis* GeneXpert (Xpert MTB/RIF Ultra) results also reported. **d.** Non-progressor contacts are grouped based on IGRA and PET-CT status after excluding clinical and radiographic outliers; IGRA^-^ PET-CT^-^ (N=9), IGRA^+^ PET-CT^-^ (N=5), IGRA^+^ PET-CT^+^ (N=6), Progressors (N=8). Each column represents an individual contact, with the latest available timepoint post-baseline shown. Where available for progressors, pre-progression and progression samples are shown (B, baseline; F, follow-up; P, progression).

### Opposing neutrophil and T cell responses in both TB patients and progressors

To further probe the molecular and cellular changes in airway cells of TB patients and their contacts, scRNA-seq analysis was performed on 894,159 total BAL leukocytes. Integrated analysis of the combined study groups revealed clusters consistent with alveolar macrophages, monocyte-derived macrophages, monocytes, conventional DCs, two clusters of neutrophils and six clusters comprising T and NK cells (**Fig. 1b; Supplementary Tables 4 and 5**). Bulk RNA-seq, scRNA-seq and flow cytometry data revealed airway immune heterogeneity amongst patients with active TB, with neutrophils dominating in around 40% of TB patients, all of whom had positive *M. tuberculosis* cultures from BAL (**Fig. 1c; Extended Data Figs. 2 and 3a-b, Supplementary Tables 1b and 5)**, while those low in neutrophils were generally high in T cells and associated signatures (**Fig. 1c; Extended Data Fig. 2 and Supplementary Table 5**). BAL from half of the progressors sampled at progression was also dominated by neutrophils, with lower T cell frequency, whilst the remaining progressors showed elevated T cells and few neutrophils **(Fig. 1d; Extended Data Fig. 3a-b; Supplementary Table 5)**. BAL of non-progressing contacts mostly contained alveolar macrophages **(Fig. 1d, Supplementary Table 5),** consistent with a largely non-inflammatory airway environment in these contacts who remained healthy. Differential abundance analysis of scRNA-seq data from IGRA^+^ PET-CT^-^ and IGRA^+^ PET-CT^+^ compared to IGRA^-^ non-progressors showed increases only within distinct T cell clusters, but no increase in neutrophils (**Fig. 2a-b**). Compared with IGRA^+^ PET-CT^+^ non-progressors, the progressor group exhibited increased abundance of neutrophils and increases across the breadth of T/NK cell clusters. In active TB patients, there was an even greater increase in neutrophils, but a less substantial increase in T/NK cell clusters, reflecting the marked dominance of neutrophils over T cells in culture-positive TB as compared to culture-negative TB patients (**Fig. 2a-b).** An increase in monocytes accompanied neutrophil accumulation during TB progression (**Fig. 2a-b**). BAL neutrophil counts by flow cytometry correlated with radiographic extent of disease and bacterial burden in active TB patients (**Fig. 2c-d**), consistent with neutrophil accumulation in more severe TB disease. Accordingly, BAL supernatants of culture-positive TB patients contained high concentrations of mediators associated with inflammation and neutrophil responses, including IL-1β, IL-1 receptor antagonist (IL-1RA), MMP9 and the neutrophil chemokine CXCL8, while the chemokines CCL24 and CXCL16 were found at similar concentrations across all groups (**Extended Data Fig. 3c-e**). It is notable that the inverse relationship observed between neutrophil and T cell frequencies in the BAL of human TB patients and progressors is consistent with the functional opposition demonstrated between these cell types in experimental mouse models^19,20^, supporting their validity for study of immunological mechanisms of TB progression and pathology.

**Fig. 2.**
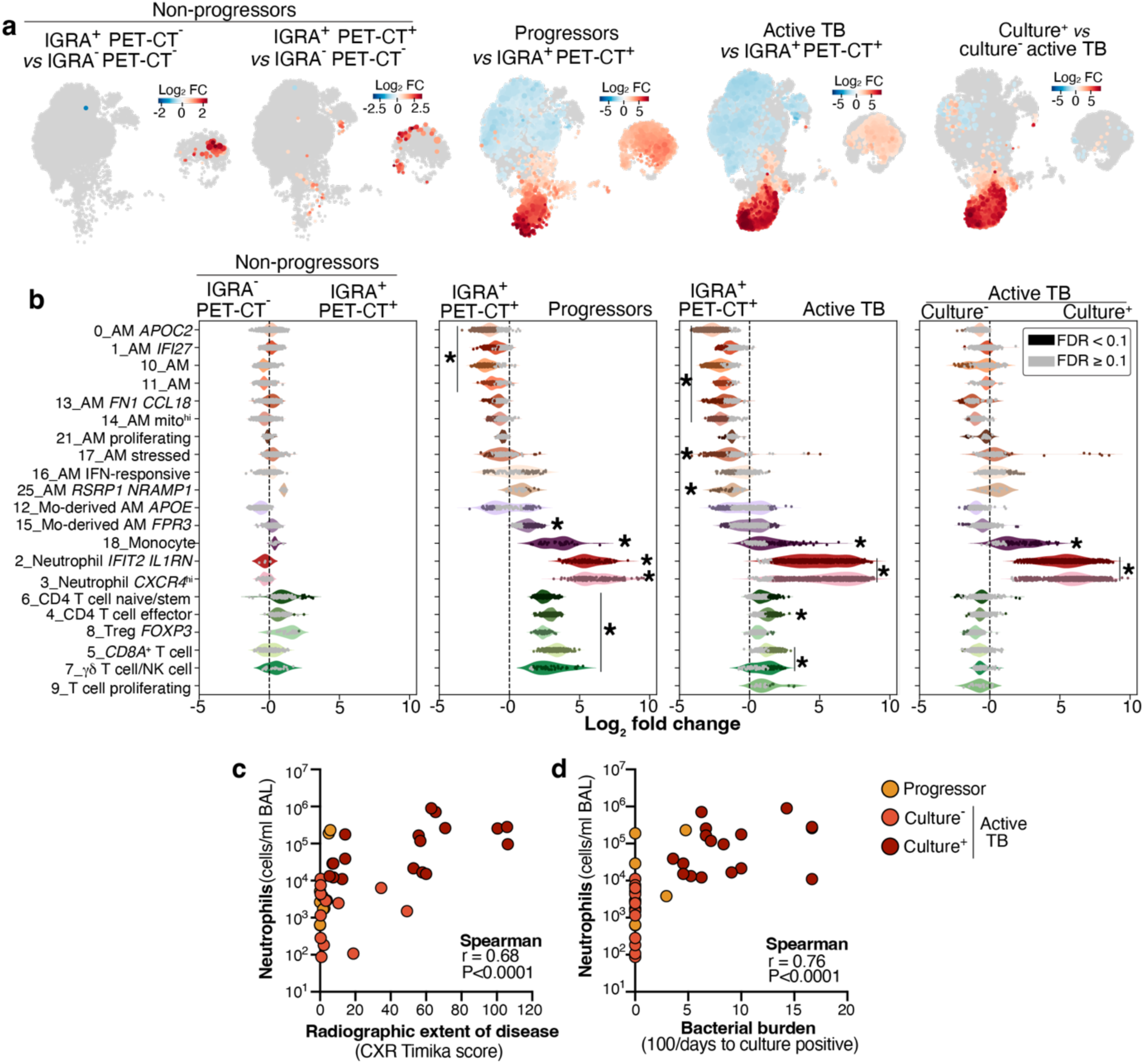
Airway neutrophils increase with TB disease progression. **a-b.** Milo differential abundance analysis of scRNA-seq cell clusters compared between the indicated groups. IGRA^-^ PET-CT^-^ (N=9), IGRA^+^ PET-CT^-^ (N=5), IGRA^+^ PET-CT^+^ (N=6), Progressors at progression (N=6); BAL culture^-^ TB (N=9) BAL culture^+^ TB (N=12). **a.** Differentially abundant cell neighborhoods are colored by Log_2_ fold change between groups and superimposed onto UMAPs. **b.** Cell neighborhoods were superimposed onto corresponding BAL leukocyte clusters as defined in **Fig. 1b**. Each dot represents one neighborhood, with fold change and FDR results for differential abundance indicated. *, marks clusters in which ≥20 % of assigned neighbourhoods were significantly differentially abundant in a consistent direction (false discovery rate, FDR <0.1) between the indicated groups. **c-d**. Neutrophil counts in progressor (N=10) and BAL culture-negative (N=13) and -positive (N=17) TB patient BAL as determined by flow cytometry, plotted per-patient against chest X ray Timika score or estimated BAL *M. tuberculosis* bacterial burden (100/days to culture positivity, with a value of 0 indicating negative BAL culture). Spearman correlation results are shown.

**Fig. 3.**
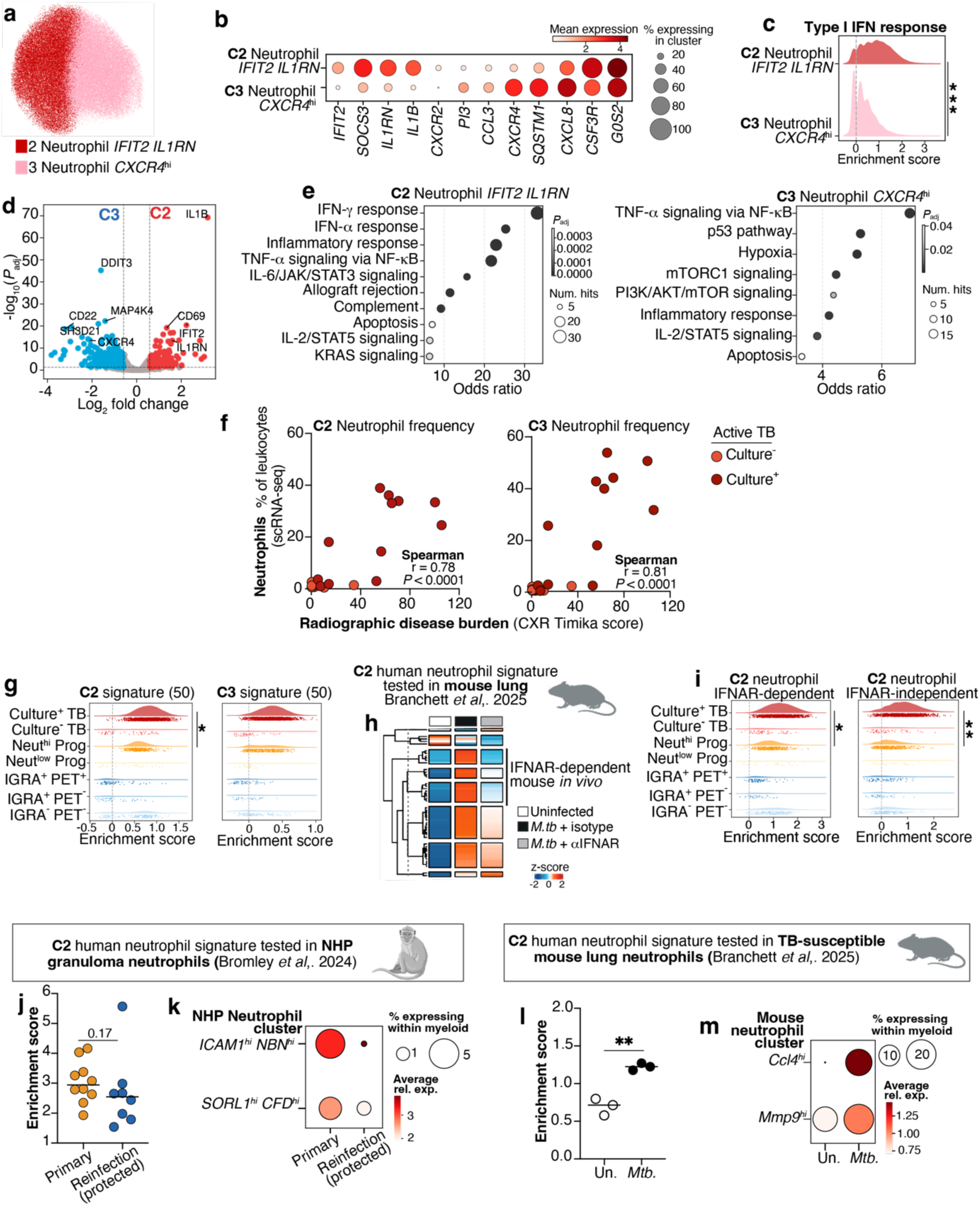
Type I IFN-dependent and -independent inflammatory neutrophil states correlate with TB progression and severity. **a.** UMAP visualization of the two neutrophil scRNA-seq clusters, cluster 2 (C2) and C3. **b.** Dot plot showing representative distinct and shared marker genes in neutrophil clusters. **c.** Density plot showing distributions of enrichment scores of a myeloid type I IFN response signature (defined in **Extended Data Fig. 4b**) in the neutrophil clusters. Statistics: Wilcoxon matched pairs signed rank test comparing mean enrichment scores per sample between C2 and C3 neutrophils from all participants with ≥20 cells in both neutrophil populations **, *P*<0.001 (N=42). **d.** Volcano plot of differentially expressed genes between neutrophil clusters across all samples. **e.** Top enriched pathways within significantly upregulated genes in neutrophil clusters. **f.** Abundance of C2 and C3 neutrophils within BAL leukocytes of TB patients plotted against chest X-ray Timika score. Spearman correlation results are shown. **g.** Distributions of enrichment scores for the top 50 differentially expressed genes (as ranked by adjusted *P* value) in C2 and C3 neutrophils across total neutrophils in each group. Dots under density plots represent individual cells **h.** Top 50 genes in C2 neutrophils were examined by k-means clustering in lung bulk RNA-seq data from mice infected with *M. tuberculosis* treated with either anti-IFNAR blocking antibody (αΙFNAR) or isotype control. Gene clusters showing IFNAR-dependency in mouse TB are highlighted. Columns in **h** show data averaged from 3-4 mice per group. **i.** Density plots as in **g** showing C2 neutrophil genes identified as IFNAR-dependent and -independent in mouse lung. **j.** Mean enrichment scores of the top 50 C2 neutrophil genes shown in total neutrophils from NHP granulomas in primary (N=10) *M. tuberculosis* infection compared to reinfection (N=8). Points show data from individual granulomas with lines at the median; statistics: Mann-Whitney test. **k.** Relative expression of C2 neutrophil genes in the two published NHP neutrophil clusters. **l**. Mean enrichment scores of the top 50 C2 neutrophil genes shown in total neutrophils per mouse from lungs of uninfected (Un., N=3) mice and at day 20 post-infection with *M. tuberculosis* (*Mtb* N=3). Statistics: unpaired t-test, **, *P*<0.01. **m.** Relative expression of C2 neutrophil genes in mouse TB neutrophil clusters. Group sizes for human scRNA-seq analysis: IGRA^-^ PET-CT^-^ (N=9), IGRA^+^ PET-CT^-^ (N=5), IGRA^+^ PET-CT^+^ (N=6), neutrophil-low progressors (N=3); neutrophil-high progressors (N=3) and BAL culture-negative (N=9) and - positive (N=12) TB patients. Statistics in **g** and **i**: Mann-Whitney test comparing mean enrichment scores per sample between BAL culture-positive TB (N=12) and neutrophil-high progressors (N=3); *, *P*<0.05, **, *P*<0.01.

### Type I IFN-dependent and independent neutrophil states increase with TB progression and severity

The two neutrophil clusters identified by scRNA-seq appeared to represent distinct inflammatory states. While cluster 3 (C3) was distinguished by the marker gene *CXCR4* and higher expression of *DDIT3* and the chemokines *CCL3* and *CXCL8,* cluster 2 (C2) was marked by high expression of *IL1B, IL1RN,* encoding IL-1RA (**Fig. 3a-b; Extended Data Fig. 4a)**, and type I IFN-inducible gene signatures (**Fig. 3c; Extended Data Fig. 4b**). IL-1β and IL-1RA proteins were also elevated in BAL of culture-positive TB patients (**Extended Data Fig. 3c-d**), in whom neutrophils were generally high (**Fig. 1c**, **Fig. 2, Extended Data Fig. 3b**).

Co-expression of type I IFN-inducible genes and *IL1B* by C2 neutrophils is notable since, while type I IFN has shown adverse effects in TB^13,14^, IL-1 signalling has been reported to be protective and functionally antagonises pathogenic type I IFN signalling in a mouse TB model^21^. Since our neutrophil C2 also expressed high levels of *IL1RN* (**Fig. 3b**), which antagonises IL-1 signalling, this may represent a feedback mechanism to control IL-1 function and untoward inflammation during chronic infection. Such feedback regulation is further supported by reports that IL-1RA is induced by type I IFN, and its deletion in TB-susceptible mice reduces type I IFN-driven TB disease^13^. *Il1b* expression and type I IFN responses have recently been reported to characterise two different hubs within the mouse NeuMap of neutrophil states across tissues and conditions^22^. It is therefore likely that neutrophil C2 in our analysis, where co-expression of *IL1B* and IFN-inducible genes was observed, represents a distinctive activation state of neutrophils in TB, a chronic intracellular bacterial infection.

The frequency of both neutrophil clusters C2 and C3 within BAL leukocytes correlated with radiographic TB disease burden (**Fig. 3f**). Both scRNA-seq clusters, as well as total neutrophil frequency by flow cytometry, were also elevated in progressing contacts with a high lung PET-CT signal (**Extended Data Fig. 4c-d**). However, while the C3 neutrophil signature genes were comparably enriched in neutrophils from progressors and TB patients, C2 neutrophil genes showed significantly greater enrichment in active TB compared to progressors (**Fig. 3g, Extended Data Fig. 4a**), suggesting that this highly inflammatory neutrophil signature increases with more advanced disease. Leveraging our published RNA-seq data of IFNαβ receptor (IFNAR) blockade in a severe mouse TB model^14,19^, the human C2 neutrophil signature genes were found to segregate into clusters of IFNAR-dependent (e.g. *IFIT2*, *IFIT3*) and -independent (e.g. *ICAM1*, *CD274*) genes in mouse lung during *M. tuberculosis* infection (**Fig. 3h**). The type I IFN-dependent and -independent C2 signatures were found to be enriched in neutrophils of neutrophil-high TB patients and progressors but both showed significantly greater enrichment in TB compared to progressors (**Fig. 3i**), suggesting that both type I IFN-dependent and -independent neutrophil activation signatures of airway neutrophils increase with more advanced TB disease.

### The human TB BAL airway neutrophil signature is recapitulated in neutrophils from lung tissue of TB disease in NHP and mice

To examine our human TB disease-associated neutrophil signature directly at the site of *M. tuberculosis* infection, we analysed published scRNA-seq data from lung granulomas of non-human primates (NHP), during either primary TB disease or a secondary *M. tuberculosis* infection where CD4 T cell-dependent protection had been achieved through a previous antibiotic treated infection^15^. Our human C2 neutrophil signature was found to be highly enriched in neutrophils from the NHP granuloma during primary infection (**Fig. 3j**). Two neutrophil subclusters identified in the NHP study both expressed this C2 signature during the primary NHP infection but were much less abundant in the granulomas of protected reinfected animals (**Fig. 3k**). This human C2 signature was also enriched in inflammatory lung in two lung neutrophil clusters in infected TB-susceptible mice, where pathogenesis is dependent on highly activated neutrophils (**Fig. 3l-m**)^14,19^. Our transcriptomic analysis of human airways therefore reveals neutrophil signatures of TB disease with type I IFN-dependent and - independent components that are consistent across species.

### Dysfunctional T cell and macrophage states in TB patients and progressing contacts with high airway neutrophils

To understand the potential impact of neutrophils on airway T-cell and macrophage responses, we compared quantitative and qualitative changes at single cell level between neutrophil-high and neutrophil-low subgroups of TB patients and progressors. Bioinformatic receptor-ligand analysis predicted extensive macrophage-macrophage and T cell-macrophage interactions in neutrophil-low TB patients and progressors, which were not apparent in the neutrophil-high state, where neutrophils dominated the predicted sending and receiving of signals (**Extended Data Fig. 5a-b**). Exclusion of neutrophil-dependent interactions from the visualisation revealed underlying T cell-macrophage and macrophage-macrophage interactions, but at a much reduced level than in the neutrophil-low state. **(Extended Data Fig. 5c).** This suggests that neutrophils may inhibit, as well as obscure, T cell-macrophage interactions, supporting observations in TB-susceptible mouse models of excessive neutrophil recruitment opposing protective T-cell responses^19,20^. Increased IL-1 and TNF pathway activity was predicted in neutrophil-high compared to neutrophil-low patients and progressors, although IL-1 pathway activity was largely predicted to the non-signalling decoy receptor, encoded by *IL1R2*, on neutrophils themselves (**Extended Data Fig. 6a-d**). Together with their high expression of IL-1RA (**Fig. 3b**), these data support a model in which type I IFN-activated neutrophils contribute to dysfunctional IL-1-IL1R1 signalling in the human TB lung. In contrast, predicted IFN-γ signalling was comparable between the neutrophil-high and -low groups (**Extended Data Fig. 6b,e**), consistent with conclusions from vaccine studies that IFN-γ alone is insufficient as a correlate of protection against TB^5^, and experimental TB model studies showing that excessive IFN-γ can contribute to immunopathology^23^. Predicted CXCL chemokine interactions in neutrophil-high TB patients and progressors were dominated by CXCL8-CXCR1/2 signals between neutrophils (**Extended Data Fig. 6b,f)**, in keeping with results in TB-susceptible mouse models, where high expression of the functional CXCL8 homologue, CXCL2, is proposed to drive swarming of neutrophils at the site of *M. tuberculosis* infection^19,24^. CXCL16-CXCR6 interactions between myeloid cells and T cells were dominant in neutrophil-low individuals, while T cell-derived CCL5 was the major predicted CCL chemokine signal and was largely preserved in the neutrophil-high state (**Extended Data Fig. 6b,g**). Collectively these data suggest that a dysregulated inflammatory response contributes to progression of TB pathology and failed bacterial control, through effects on protective macrophages and T cells, which we investigated further.

Flow cytometry analysis revealed that surface co-expression of chemokine receptors CXCR3 and CCR6 by CD45RA^-^ CCR7^-^ CD4 effector/effector memory (T_eff/em_) cells was almost completely absent in neutrophil-high TB patients and progressors (**Fig. 4a,c**). CXCR3^+^CCR6^+^ CD4 T_eff/em_ frequency inversely correlated with radiographic TB disease burden (**Fig. 4b**), consistent with T cell dysfunction and failed immune control of *M. tuberculosis* associating with the neutrophil-high state. Our findings support reports that CXCR3^+^CCR6^+^ marks a putative protective phenotype of *M. tuberculosis* antigen-specific CD4 T cells in human blood^25^ and NHP BAL^26^ in latent TB. In contrast to expression of CXCR3^+^CCR6^+^, expression of the activation marker HLA-DR on CD4 T_eff/em_ cells showed no correlation with the extent of neutrophilic airway inflammation **(Extended Data Fig. 7a-b**). To better understand the dysfunctional T cells associated with high levels of neutrophils, differential gene expression analysis of T cell scRNA-seq clusters was performed. Compared with neutrophil-low groups, higher levels of coinhibitory receptors, cytotoxic mediators and genes related to apoptotic and inflammatory cell death were observed in CD4 T effector and *CD8A*^+^ T cell clusters from neutrophil-high TB patients and progressors (**Fig. 4d; Extended Data Fig. 7c**). Moreover, activity of the TRAIL and TWEAK pathways of extrinsic apoptosis was predicted in the neutrophil-high but not neutrophil-low groups (**Fig. 4e-f**). These data suggest that chronic activation, exhaustion and cell death in airway T cells likely contribute to failed immune control of *M. tuberculosis* accompanying excessive neutrophil accumulation.

**Fig. 4.**
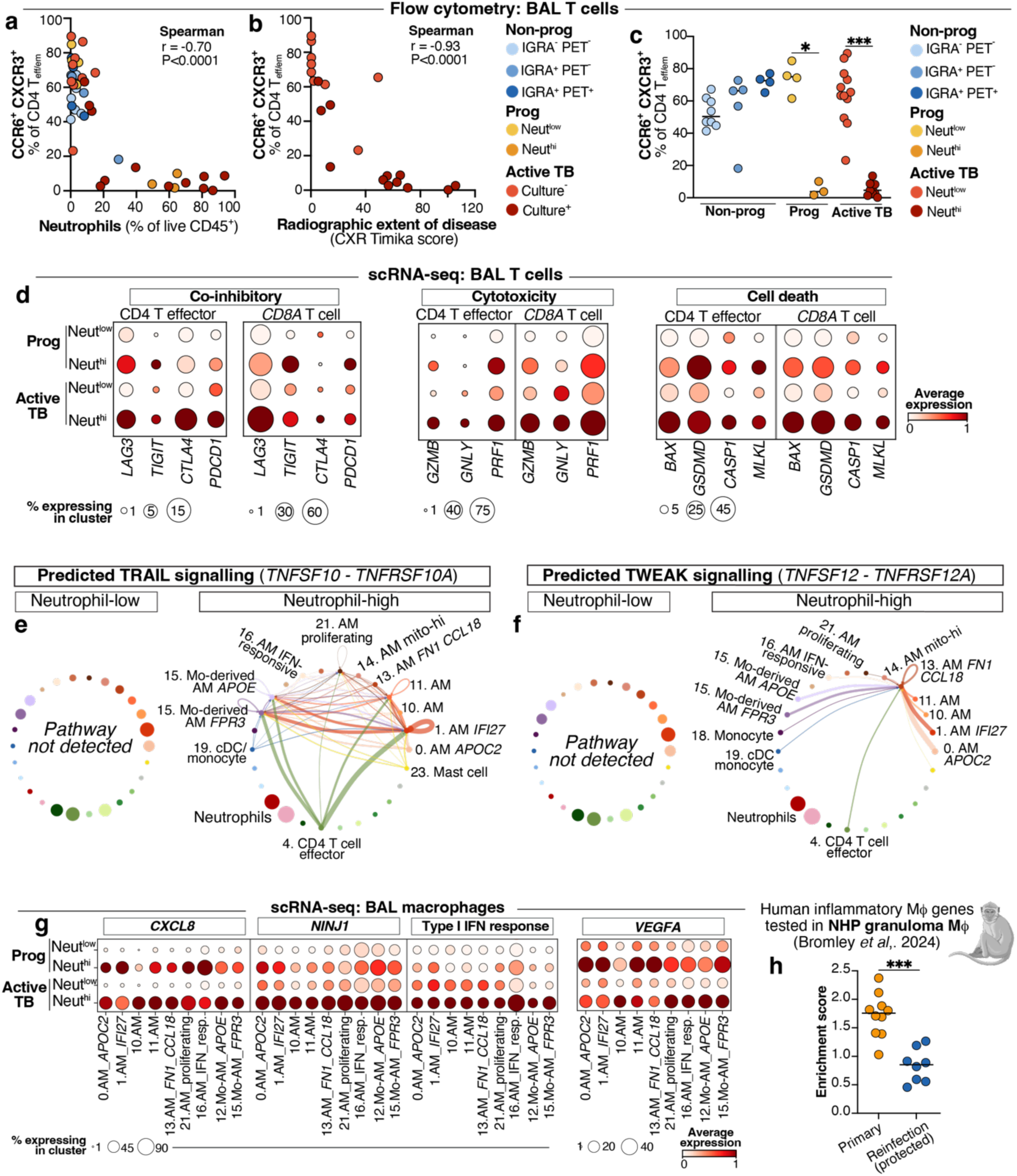
Neutrophil accumulation marks a dysfunctional airway immune cell state in TB patients and progressing contacts. **a-b.** Frequencies of CCR6^+^ CXCR3^+^ CD4 T_eff/em_ cells within total CD4 T_eff/em_, as determined by flow cytometry in IGRA^-^ PET-CT^-^ (N=8), IGRA^+^ PET-CT^-^ (N=5), IGRA^+^ PET-CT^+^ (N=4), neutrophil-low progressors (N=4), neutrophil-high progressors (N=3) and BAL culture-negative (N=9) and -positive (N=12) TB patients. Neutrophil-low and -high states were defined as < or ≥ 15% neutrophils, respectively, as a percentage of live, CD45^+^ BAL leukocytes by flow cytometry. Data are plotted against proportion of neutrophils in BAL and chest X-ray Timika scores, and Spearman correlation results are shown. **c.** Flow cytometry as in **a-b** shown across study groups, with active TB patients split into neutrophil-low (N=12) and -high (N=9) sub-groups. Points show individual participants with lines at the median. Statistics: Kruskal-Wallis test with Dunn’s *post hoc* test; *, *P*<0.05; ***, *P*<0.001. **d.** Expression of representative genes increased in airway T cells from neutrophil-high (N=11) compared to neutrophil-low (N=16) active TB patients and progressors. **e-f.** CellChat analysis showing predicted activity of TRAIL and TWEAK pathways between leukocyte populations in BAL. Line colour indicates the predicted signal-sending cell cluster and line thickness indicates predicted interaction strength. **g.** Expression of representative genes and gene sets increased in airway macrophages from neutrophil-high compared to neutrophil-low active TB patients and progressors. Type I IFN response refers to the gene expression signature defined in **Extended Data Fig. 4b**. Data shown in **d-g** are averaged from all cells from all samples per group: neutrophil-low progressors (N=3); neutrophil-high progressors (N=3) and neutrophil-low TB (N=13) and neutrophil-high (N=8) TB patients **h.** Mean enrichment scores of a 50 gene signature of human airway inflammatory macrophages (see **Methods** and **Extended Data Fig. 7d-e**) shown in total macrophages from NHP granulomas in primary (N=10) *M. tuberculosis* infection compared to reinfection (N=8). Points show data from individual granulomas with lines at the mean; statistics: unpaired t-test, ***, *P*<0.001.

Dysregulated inflammatory gene expression was also observed across BAL macrophages in neutrophil-high patients and progressors, enriched for functions such as chemotaxis, inflammatory cell death and VEGF signalling (**Fig. 4g; Extended Data Fig. 7d-f**). Macrophage expression of *CXCL8* was increased in the neutrophil-high state, suggesting that macrophages may contribute to continued neutrophil recruitment, although *CXCL8*-expressing neutrophils far outnumbered macrophages in these patients (**Fig. 4g; Extended Data Fig. 6b,f**). Additionally, increased macrophage type I IFN-inducible gene expression was observed in the neutrophil-high state and more pronounced in active TB than progressors (**Fig. 4g**), mirroring results in neutrophils (**Fig. 3i**). A combined human airway macrophage transcriptional signature of failed immunity in neutrophil-high TB was highly enriched in granuloma macrophages of *M. tuberculosis*-infected macaques and diminished in the context of protective immunity from prior infection^15^ (**Fig. 4h; Extended Data Fig. 7e,g**). Conversely, expression of genes related to core alveolar macrophage functions such as lipid metabolism, oxidative phosphorylation and organelle homeostasis decreased across airway macrophage populations of progressing contacts compared to non-progressors (**Extended Data Fig. 8a-c**) and this gene signature was enriched in alveolar macrophage-like cells present in granulomas of immune-protected macaques, supporting cross-species consistency of macrophage transcriptional responses to *M. tuberculosis* infection (**Extended Data Fig. 8d**).

### Cross-species T cell signatures of *M. tuberculosis* control revealed in airways of human TB contacts

We further investigated our datasets for evidence of active protective immune responses in non-progressor contacts. Reasoning that IGRA^+^ non-progressors with a positive thoracic PET-CT represent those mounting T-cell dependent responses to current *M. tuberculosis* infection, we compared effector T cells in this group to the other non-progressors in our study. The frequency of HLA-DR^+^ activated T cells, particularly those also lacking PD-1 surface expression, was increased among BAL T_eff/em_ in IGRA^+^PET-CT^+^ compared to IGRA^-^ PET-CT^-^non-progressors (**Fig. 5a; Extended Data Fig. 9a**). Abundance of these T cells was greatest in those IGRA^+^ non-progressors with positive thoracic LN, but not lung, PET-CT (**Fig. 5b; Extended Data Fig. 8a**), suggesting that a PET-CT signal in the LN of non-progressors may be associated with a protective airway T cell response. In support of this, we show that a protective T1-T17 signature of *M. tuberculosis* control from NHP granulomas^12^ was enriched in our scRNA-seq CD4 T-effector population from human BAL of IGRA^+^ LN PET-CT^+^ non-progressors (**Fig. 5c; Extended Data Fig. 9c**).

**Fig. 5.**
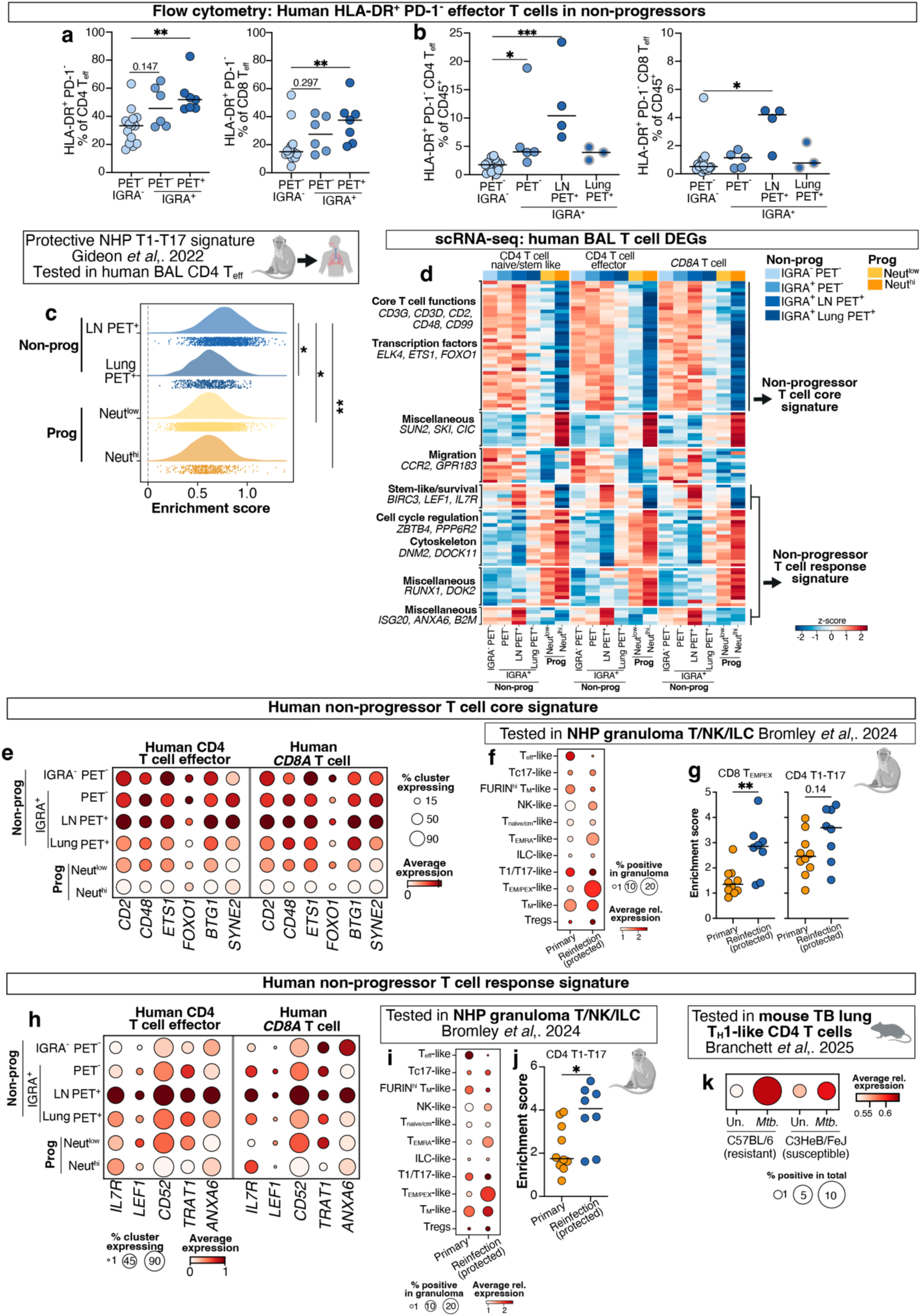
Cross-species T cell signatures of pulmonary *M. tuberculosis* infection control. **a-b.** Flow cytometry data from BAL of non-progressor contacts showing HLA-DR^+^ PD-1^-^ CD4 and CD8 T_eff/em_ cells either as a percentage of **a.** the parent T_eff/em_ population or **b.** total live CD45^+^ leukocytes. Points show individual contacts with lines at the median; IGRA^-^ PET-CT^-^(N=15), IGRA^+^PET-CT^-^ (N=5), IGRA^+^ LN PET-CT^+^(N=4), IGRA^+^ lung PET-CT^+^ (N=3). Statistics: Kruskal-Wallis test with Dunn’s *post hoc* test; *, *P*<0.05; **. *P*<0.01; ***, *P*<0.001. **c.** Distributions of enrichment scores for the top 50 signature genes of the T1-T17 T cell population associated with protection in NHP granulomas in the CD4 T effector cluster from human BAL. Combined data from N=3 individuals per group are shown. Dots under density plots represent individual cells. Statistics: Kruskal-Wallis test with Dunn’s post hoc test comparing mean enrichment scores from each participant (N=3); *, *P*<0.05, **, *P*<0.01. **d.** Heatmap showing k-means clustering of shared differentially expressed genes across major scRNA-seq T cell clusters combined from comparisons between progressor and non-progressor sub-groups (detailed in **Supplementary Table 7**); IGRA^-^ PET-CT^-^ (N=9), IGRA^+^ PET-CT^-^ (N=5), IGRA^+^ LN PET-CT^+^ (N=3), IGRA^+^ lung PET-CT^+^ (N=3), neutrophil-low progressors (N=3) and neutrophil-high progressors (N=3). **e,h.** Expression of representative genes from non-progressor-associated T cell signatures across TB contact groups. **d,e,h** show averaged data across all samples per group. **f-g, i-j.** Human BAL non-progressor T cell signatures were examined in lung granuloma T/NK cell scRNA-seq data from NHP and compared between those from primary *M. tuberculosis i*nfection (N=10) and in the context of immune protection during reinfection (N=8). **f,i.** Expression of the indicated signatures in in the indicated NHP granuloma scRNA-seq clusters, averaged across all granulomas in each group. **g, j.** Mean enrichment scores of human non-progressor T cell signatures in the indicated populations from NHP granulomas. Points represent data from individual granulomas with lines at the median; statistics: Mann-Whitney t-test, *, *P*<0.05; **, *P*<0.01. **k.** Average expression of the human BAL non-progressor T cell response signature in T_H_1-like CD4 T cells in mouse lung in uninfected (un.) mice and at 20 days post-infection (*Mtb*). Averaged data from N=3 mice per group are shown.

To further explore potentially protective T cell responses in the airways of non-progressing contacts, we performed differential expression analysis in our human scRNA-seq dataset, identifying common patterns of differentially expressed genes among the major T cell clusters between progressor and non-progressor sub-groups (**Fig. 5d; Extended Data Fig. 9d**). A non-progressor T cell core signature was found to be expressed in IGRA^-^, IGRA^+^PET-CT^-^ and IGRA^+^ LN PET-CT^+^ non progressors and reduced in lung PET-CT^+^ non-progressors and TB progressors (**Fig. 5d; Extended Data Fig. 9e-f**). This signature included cell-cell adhesion genes *CD2* and *CD48*, the transcriptional regulator *ETS1* and the T cell-quiescence factors *FOXO1^27^* and *BTG1^28^* (**Fig. 5d-e**), suggestive of a restrained activation state that is diminished with progression to TB. This quiescence signature was observed in the *CD8A*^+^ T cell and CD4 T effector clusters and not restricted to the CD4 naïve/stem-like cluster **(Fig. 5d-e)**, suggestive of a common functional state across conventional T cell subsets in non-progressors, rather than simply a relative reduction in naïve T cells in BAL of contacts as they progressed to TB. Indeed, our flow cytometry analysis showed that the vast majority of airway CD4 T cells across groups had a CD45RA^-^ CCR7^-^ T_eff/em_ phenotype (**Extended Data Fig. 9b**). We determined that this human non-progressor T cell core signature was also increased in the granuloma T cells of NHP that had been immune protected as compared to primary infection and disease^15^ (**Fig. 5f)**. This was most prominently observed in the NHP *GZMK*^hi^ CD8 T effector memory/precursor exhausted (T_EM/PEX_)-like cells (**Fig. 5f-g**), a population that was more abundant in granulomas in the context of immune protection in the published NHP study^15^.

A second T cell response signature was strikingly increased in non-progressors with positive LN but not lung PET-CT signal and was markedly reduced in TB progressors, consistent with a potentially protective T cell state (**Fig. 5d; Extended Data Fig. 9g-h**). This signature of LN PET-CT^+^ IGRA^+^ non-progressors distinguished this group from PET-CT^-^ IGRA^+^ and IGRA^-^non-progressors (**Fig. 5d; Extended Data Fig. 9g-h**) suggesting that this signature potentially discriminates the immune response to a recent *M. tuberculosis* infection from a cleared or past infection. This non-progressor T cell response signature included *IL7R* and *LEF1*, both associated with a stem-like T cell state^29,30^. Accompanying this stem-like profile was expression of genes implicated in regulation of T cell receptor signalling and membrane dynamics, *CD52^31,32^*, *TRAT1*^33^ and *ANXA6*^34^(**Fig. 5d,h**). Collectively, our findings in human airways of TB contacts who controlled infection suggest that a restrained, stem-like activation state is maintained to ensure durable protection from progression to TB disease.

We then tested our human airway non-progressor stem-like T cell signature response in scRNA-seq data from T cells within granulomas of immune protected NHP. Expression of this signature was increased in T cells from immune protected NHP as compared to primary infection and disease, particularly in the abundant CD8 T_EM/PEX_-like cluster, as well as the smaller CD4 T1-T17 effector cluster (**Fig. 5i-j**). We additionally observed this signature in T_H_1-like CD4 T cells in lungs of relatively TB-resistant C57BL/6 mice early in infection, at higher frequency than in TB-susceptible C3HeB/FeJ mice that fail to control *M. tuberculosis* infection^19^ (**Fig. 5k**). A stem-like transcriptional signature, including *TCF7*, *LEF1* and *SELL*, was also enriched in IGRA^+^ non-progressors and decreased with progression within a small cluster of γδ/NK/ILC cells identified in our scRNA-seq analysis (**Extended Data Fig. 10a-h**). In keeping with this, NK cells have been associated with controlled *M. tuberculosis* infection in human blood^11,35^ and NHP lung^11^. Our human stem-like signature of innate-like lymphocytes showed greater enrichment in NHP granulomas from immune protected animals, compared to primary infection (**Extended Data Fig. 10d-e**). Thus, we show that stem-like transcriptional features within effector T cells and innate lymphocytes, which are associated with control of *M. tuberculosis* at the site of pulmonary infection in airways of non-progressing human TB contacts, are recapitulated in lung tissue of animal models representing improved TB control.

## Discussion

While most individuals infected with *M. tuberculosis* control the infection and remain asymptomatic, the local early immune factors that dictate protection or disease progression have not been defined. We have charted the airway immune response at single cell resolution in an extensively characterised and prospectively observed cohort of household TB contacts, revealing early immune changes at the site of *M. tuberculosis* infection that reflect protection in contacts who remained clinically healthy and radiologically stable by PET-CT and, conversely, inflammatory signatures of failed immunity in individuals who progressed to TB. Using unbiased analysis of bulk and scRNA-seq of BAL samples, we here report an inverse relationship between neutrophils and T cells in TB patients and contacts that progress to TB, with T cells in progressors showing signatures of exhaustion, cytotoxicity and cell death. T cell signatures of protection in contacts who remained healthy were dominated by regulation, quiescence and a stem-like profile. These human airway immune signatures of protection and progression were recapitulated in lung tissue from NHP and mouse TB models with controlled infection or disease, respectively.

In-depth analysis of scRNA-seq data from airway neutrophils revealed two distinct transcriptional states in culture-positive TB patients and a subset of TB progressors, with both type I IFN-dependent and type I IFN-independent inflammatory signatures associating with more advanced disease. Lung neutrophil accumulation is a common endpoint of failed immune control of *M. tuberculosis* in mouse models^14,24,36^ and increased neutrophil numbers in BAL are a feature of more advanced active TB^37^. Our current findings suggest that neutrophils may drive TB progression as well as advanced disease. While a blood neutrophil type I IFN-inducible signature in humans has been associated with more advanced TB disease^9,10,16^, IFNAR blockade only partially protected genetically susceptible mice from severe TB^14,19^, supporting the role of type I IFN-independent mechanisms of neutrophil-driven TB progression, as supported by our current human TB contact study. Airway neutrophil accumulation in human TB patients and progressors was accompanied by profound transcriptional and phenotypic changes in airway T cells, suggestive of chronic activation and impaired function, along with widespread inflammatory changes in macrophage populations. These findings in human airways support mechanistic studies in mouse TB models, where neutrophil accumulation opposes protective immune interactions in the lung^19,20^. Our findings that inflammatory neutrophil and macrophage signatures of TB disease and progression in human airways are captured in granuloma scRNA-seq datasets from NHP during primary infection and disease and reduced in the context of immune protection^15^ indicates that BAL samples can reflect pathogenic immune responses present in the granuloma.

Determinants of an effective protective T cell response to *M. tuberculosis* are not well understood but are likely to depend upon temporal and spatial factors, such as durability of responses and ability of T cells to access and interact with infected macrophages^6,7^. Our findings that airway CD4 and CD8 T cells from IGRA^+^ LN PET-CT^+^ non-progressors were enriched for T cell signatures of stem-like memory T cell states suggests their role in active immune protection. This is supported by numerous reports of stem-like memory CD8 T cells providing durable protective memory responses to tumours and chronic viral infections^29,30,38-41^. Our findings of airway stem-like memory T cell states in CD4 T cells of non-progressing TB contacts are in keeping with reported detection by flow cytometry of antigen-specific T stem cell memory CD4^+^ T cells in the blood of BCG-vaccinated individuals which correlated with a long-term T cell proliferative response^42^. Additionally, a stem-like transcriptional profile has been reported in *M. tuberculosis* antigen-specific CD4 T cells in the blood from individuals who control infection, with *LEF1* and *IL7R* specifically increased in IGRA^+^, compared to persistently IGRA^-^ individuals termed resistors^43^. Our demonstration of a stem-like airway T cell signature, specifically in IGRA^+^ LN PET-CT^+^ non-progressors, suggests that it is part of a protective response in the airways of recent TB household contacts, reflecting current or recent controlled *M. tuberculosis* infection. Our comparative studies demonstrating enrichment of our human BAL T cell signatures of stem-ness and restrained activation in the context of adaptive immune protection in NHP granuloma T cells^15^, supports an association of these transcriptional states with *M. tuberculosis* control.

Our study of recent human TB household contacts in a low TB burden setting reveals early immune signatures at the site of *M. tuberculosis* infection associated with infection control or disease progression. Recapitulation of these human signatures in datasets from lungs of experimental animal TB models exhibiting controlled infection or disease strongly supports their involvement in dictating infection outcome. While signatures of disease progression provide targets for further study in the context of host-directed therapy, our stem-like T cell signatures in contacts demonstrating active immune control will be of use to inform strategies for improved vaccines against TB.

## Methods

### Participant recruitment and clinical study design

This was a 24-month prospective observational cohort study of pulmonary TB patients and recent household TB contacts, recruited between September 2021 and April 2024 at University Hospitals of Leicester NHS Trust, UK (ISRCTN1798557). The study was approved by the Research and Ethics Committee (REC) for East Midlands - Nottingham 1, Nottingham, UK (REC 21/EM/0139). Eligible participants were immunocompetent adults aged ≥16 years with suspected TB or recent household contacts of pulmonary TB. Exclusion criteria were significant immunosuppression caused by immunodeficiency disorders or use of T-cell specific or generalised immunosuppressive therapy, including oral corticosteroids at any dose; history of receiving TB preventive therapy (TPT) in the preceding 5 years, or previous treatment for active TB in the preceding 2 years; and pregnancy or lactation.

Thirty-two adults with pulmonary TB were recruited to the study and underwent BAL sampling. All participants had compatible clinical signs and symptoms and typical imaging features of active TB disease. In twenty-five participants, microbiological confirmation was achieved by culture and/or Xpert-MTB RIF Ultra testing of either respiratory tract samples (N=22), or from non-respiratory sites (N=3) in cases with multi-compartmental involvement. In seven participants without microbiological confirmation, the diagnosis was supported by histology and/or verified clinical and radiological response to a full course of anti-tubercular treatment. Two BAL samples from microbiologically confirmed TB patients were excluded from all analysis due to excessive blood contamination in one, and a subsequent diagnosis of chronic lymphocytic leukaemia in the other, leaving 30 samples for analysis. TB patients were further grouped based on detection of positive *M. tuberculosis* culture in BAL (**Supplementary Table 1b**).

Asymptomatic household pulmonary TB contacts were recruited soon after index case notification. Consenting participants completed a symptom questionnaire and underwent chest radiography (CXR), to exclude active TB. IGRA testing was performed with QuantiFERON-TB GOLD Plus and among IGRA-positive participants, those that declined TPT were retained in the study. All participants were offered bronchoscopy after PET-CT at baseline and after 3-months (median time after index case notification to recruitment was 23 days). Thereafter, all IGRA-positive participants were actively monitored for evidence of TB progression at 3-monthly intervals with symptom questionnaire and CXR until 24 months; IGRA-negative participants were passively followed with instructions to contact the study team if any symptoms develop (UK standard of care). At the end of 24 months, participants were contacted to verify they had remained well. A total of 157 contacts were recruited; four participants withdrew, 53 declined bronchoscopy, and 34 were not offered bronchoscopy (**Extended Data Fig. 1a**). In total, 66 pulmonary TB contacts provided BAL. Participants were grouped for analysis based on clinical progression, PET-CT features, and IGRA results over the study period.

Eleven progression events occurred over the follow-up period (**Extended Data Fig. 1a-b and Supplementary Table 1c**). Progression events were defined by the clinical decision to treat for TB disease and classified into three phenotypes: i. Clinical TB; ii. Asymptomatic TB; or iii. PET-CT–based progression. Clinical and asymptomatic TB were defined by WHO criteria. For clinical TB, this was new symptoms and/or signs of active disease, supported by compatible radiological and/or microbiological and/or histological evidence of TB. For asymptomatic TB, this was chest radiographic and/or microbiological evidence of TB in the absence of any symptoms or signs. In one case (participant #75), a progression event was defined by significant interval PET-CT progression in the absence of visible CXR changes that was clinically attributable to TB and which raised sufficient clinical (high risk if left alone) and public health concern (risk of transmission) to arrange invasive investigation. Progression was confirmed by microbiological evidence and compatible histology obtained from the presumed site of disease. BAL was obtained from 10 progressors at the point of progression. In one case (#222), bronchoscopy was not performed as progression was characterised by evolving pleural pathology in the absence of any pulmonary involvement. TB in this case was confirmed by pleural biopsy (**Supplementary Table 1c, Supplementary Fig. 1**).

Demographic details for each study subject are included in **Supplementary Table 1a-b**.

### Sample inclusions and exclusions

Current tobacco smokers were excluded from analysis of non-progressor contact groups (two IGRA^-^ PET-CT^-^, three IGRA^+^ PET-CT^-^ and three IGRA^+^ PET-CT^+^), due to potential confounding effects of smoking on the subtle immune signatures expected in non-progressors. 10 of 11 progressors and 25 of 30 active TB patients were non-smokers and no exclusions were made from these groups. Of the remaining 30 IGRA^-^ non-progressors sampled by bronchoscopy, 11 were excluded from the IGRA^-^ study group for analysis as clinical outliers due to i) respiratory symptoms close to the time of sampling attributable to a community acquired infection, with or without positive BAL bacterial (non-mycobacterial) culture or viral PCR (N=6); ii) without respiratory symptoms but having a positive BAL bacterial culture or viral PCR accompanied by elevated BAL neutrophils (N=2); iii) declined PET-CT, preventing categorisation (N=1); or iv) positive PET-CT findings of unknown significance (N=2). One IGRA^+^ non-progressor was excluded from analysis due to presenting with a transient high PET-CT signal in the lung that quickly resolved (#182). Two further IGRA^+^ PET-CT^+^ non-progressors were excluded due to lung PET signal consistent with scarring and residual inflammation from past lung disease that prevented confident PET-CT based categorisation of these subjects (#048 and #233). Unless otherwise stated, data shown in figures represent the latest available BAL sample post-recruitment from the indicated contact.

No flow cytometry was performed on the follow-up BAL samples from two IGRA^+^ PET-CT^+^ contacts (#140, #141) because of unexpected problems with equipment. One IGRA^-^ PET-CT^-^non-progressor contact (#154) was not included in flow cytometry analysis of T cells as we observed no binding of the anti-human CD4 flow cytometry antibody. This same phenomenon was observed in two further contacts excluded for other reasons (#097 and #098) and all three of these contacts were of African descent. In other cases, certain analyses were not performed for certain samples due to either insufficient cell number or samples being obtained early in the study prior to establishment of fixed cell scRNA-seq and T cell-focused flow cytometry methods. A full list of samples and data acquired per participant is provided in **Supplementary Tables 1a-b**.

### Positron Emission Tomography and Computed Tomography (PET-CT) scans and analysis

^18^F-FDG PET-CT scans were performed according to local procedure guidelines at the Leicester PET-CT Centre (Alliance Medical Ltd.). All participants were required to fast for 6 hours prior to the scanning. Female participants of childbearing age were offered pregnancy testing before PET-CT. Participants received an intravenous injection of ^8^F-FDG at a dose of 3.5 MBq/kg +/- 10% up to a maximum of 400 MBq. Imaging was performed from the skull vertex to the proximal third of the femur using the same PET-CT scanner (GE Discovery PET-CT 710, Alliance Medical Ltd) 1 hour after radiotracer administration. Images were reconstructed using a Bayesian penalized likelihood reconstruction algorithm (Q.Clear, GE Healthcare, Chicago, USA). PET-CT scans were independently quantified by two readers. The quantification method has been described previously^44^. Briefly, anonymised PET-CT DICOM images were uploaded to 3D Slicer version 5.4.0 (https://www.slicer.org). Normal physiological reference was defined as liver mean standardised uptake value (SUV) plus three standard deviations^45^. Scans were considered positive if FDG uptake exceeded the physiological reference in intrathoracic lymph nodes and/or if focal lung parenchymal uptake was greater than background lung activity. Extracted PET metrics included maximum SUV (SUV_max_), mean SUV (SUV_mean_), metabolic volume (MV), and total lesion glycolysis (TLG, calculated by SUV_mean_ x MV). Where values differed by more than 10%, they were reviewed and re-extracted to minimise variability. Final values represent the means of scores from two independent reviewers.

### Bronchoscopy

Bronchoscopies with BAL were performed between September 2021 and October 2025 by a trained respiratory physician at University Hospitals of Leicester NHS Trust. The procedure was conducted under conscious sedation according to the British Thoracic Society guidelines for diagnostic flexible bronchoscopy^46^. Participants were required to fast for six hours prior to the procedure. The flexible bronchoscope was introduced either via nasal cavity or orally through a plastic mouth guard. Sampling was directed to target lobes and lung segments identified by chest X-ray and/or PET-CT imaging. BAL was performed by instilling and aspirating 20 ml aliquots of warmed 0.9% sodium chloride solution, up to a maximum of 250 ml, with the aim of recovering ≥ 60 ml of fluid (actual recovery range 37 to 107 ml, median 66 ml). In all participants, the first 5-10 ml of BAL fluid was sent for clinical testing including routine and mycobacterial culture and viral PCR. In participants with suspected TB, a proportion of the sample was additionally sent for cytological evaluation and tested with Xpert MTB/RIF Ultra.

### Bronchoalveolar lavage processing

All sample processing, flow cytometry staining and preservation of samples for subsequent analyses was performed on the day of bronchoscopy. BAL samples were transported on gel cold packs to the Francis Crick Institute, London, UK, with a median transit time of 2 hours and 50 minutes (range 1 hour and 45 minutes to 4 hours and 4 minutes) before initiation of processing. Samples were filtered to remove mucus and debris by pipetting through 70 um cell strainers, before pelleting by centrifugation at 500 x G for 7 minutes at 4°C and transferring supernatants in aliquots to storage at -80°C for subsequent processing. All cell pellets were briefly suspended in ammonium chloride potassium (ACK) lysing buffer (Thermo Fisher Scientific) to lyse erythrocytes, before washing with RPMI 1640 supplemented with 10% foetal bovine serum and 5% penicillin and streptomycin (all Thermo Fisher Scientific) and centrifuging at 400 x G for 5 minutes at 4°C and suspending in supplemented RPMI. Numbers of cells in single cell suspensions were counted in FastRead disposable haemocytometers (Immune Systems Ltd), with viability assessed by Trypan blue exclusion. Median viability was 82.6% (range 57.7 to 100%).

### RNA extraction from BAL cell pellets

Cell suspensions containing 0.5 x 10^6^ live cells were pelleted by centrifuging at 400 x G for 5 minutes at 4°C and pellets suspended in TRI reagent (Sigma-Aldrich). Samples were vortexed at high speed for around 20 seconds before storage at -80°C until RNA extraction. RNA extraction was performed using the DirectZol RNA Microprep kit (Zymo Technologies), as per manufacturer’s instructions, including on-column DNase I digestion.

### Bulk RNA sequencing

Extracted RNA from whole BAL cell pellets was subjected to Agilent TapeStation quality control analysis and RNA integrity (RIN) values determine to be ≥7.4 and ≤9.9, except for one sample (T219_1, from active TB patient #219), which had a RIN value of 4.1. 200 ng of total RNA was used for library preparation using the RNA library prep kit with Polaris ribosomal RNA depletion (Watchmaker Genomics). Approximately 2.5 x10^7^ 100 base pair paired end reads were obtained per sample using an Illumina NovaSeq X sequencer.

Bulk RNA-seq data were processed and aligned to the GRCh38 genome using the default nf-core/rnaseq pipeline v3.16.1, built with Nextflow. Briefly, this runs Trim Galore, STAR, and Salmon for trimming, alignment, and transcript count quantification, among additional quality controls. An in-depth description of the pipeline can be found at https://nf-co.re/rnaseq/3.16.1/. Normalised transcripts per million (TPM) data were then filtered for highly variable genes, defined as those with greater than two-fold deviation from the median in more than 10% of samples. Hierarchical cluster analysis using Wards clustering was performed to assess major sources of variation across the dataset and led to the identification of epithelial-associated gene sets, associated with variable levels of contaminating lung epithelial material in BAL pellets (**Extended Data Fig. 2a**), which were removed from downstream analysis. This analysis revealed a small cluster of genes with increased expression in BAL of current tobacco smokers (**Extended Data Fig. 2a**). Differential gene expression was then performed using DESeq2 v1.42.1^47^. Expression values were normalised by vst transformation and Wald tests implemented to test for differential expression between specific groups, adjusted for sex. Genes were determined to be differentially expressed with a fold change of ≥1.5 and Benjamini–Hochberg-adjusted *P* value of <0.05. Differentially expressed genes were clustered based on their normalised expression profile across all groups, using k-means clustering. Optimal k was determined by iterating through a k from 5 to 15 and manually selecting the number of clusters that best captured the observed expression pattern.

### Single cell RNA sequencing

Cell pellets containing 0.5-1 x 10^6^ live cells were fixed and permeabilized using the Chromium Next GEM Single Cell Fixed RNA Sample Preparation Kit (10X Genomics), as per manufacturer’s instructions. Briefly, cell pellets were suspended in 0.04% Ultrapure BSA (Thermo Fisher Scientific) in 1X PBS (Corning), then centrifuged at 400 x G for 5 minutes at 4°C, before suspending in 10X Genomics Fixation and Permeabilization buffer, supplemented with 4% molecular biology grade formaldehyde, and fixing at 4 °C for 22 hours. Fixed cells were pelleted by centrifugation at 900 x G for 5 minutes, suspended in 10X Genomics Quench buffer and supplemented with 10X Genomics Enhancer solution and 50% molecular biology grade glycerol (Sigma-Aldrich), to a final glycerol concentration of 10%, before storing at - 80°C until subsequent batch processing. Sample were stored for between 7 and 574 days before running scRNA-seq. Fixed samples were processed in pooled scRNA-seq runs of up to 10 samples using the Chromium Fixed RNA Profiling Kits for Multiplexed Samples, splitting each sample over one to three barcoded probe sets, to maximise cells recovered per sample where input cell number was sufficient for splitting over multiple barcoded probe sets.

### Single cell RNA-sequencing analysis

Cell Ranger v7.0.1 raw matrices were input into CellBender v0.3.2 to model ambient RNA and remove contaminating transcripts^48^. The expected number of cells per sample was set to recover cell numbers comparable to original input. Filtered matrices from CellBender were then processed in Python v3.11.13 using scanpy v1.11.4^49^. Cells were filtered using a permissive lower cutoff of 100 total UMI counts or 100 unique genes after ambient RNA removal, and a mitochondrial transcript fraction greater than 5%, based on iterative quality control checks to retain populations with low RNA content such as neutrophils. Scrublet^50^ was then used to identify likely doublet cells. Cells marked as doublets were not excluded from the initial analysis but were instead used to identify doublet-enriched clusters downstream. Log normalisation and principal component (PC) analysis were performed using scanpy functions sc.pp.normalize_total, sc.pp.log1p and sc.tl.pca.

An integrated manifold was generated using the scanpy Harmony implementation^51^ with key set to ‘sample’, using the top 2000 variable features and first 50 PCs. The Harmony-corrected PCs were then used to generate a nearest-neighbour graph and uniform manifold approximation and projection (UMAP) embedding.

Based on the nearest-neighbour graph, single cells were annotated according to similarity to reference populations in published human lung and immune cell scRNA-seq datasets^52,53^ using CellTypist^53^. Iterative Leiden clustering to delineate cell types and recover cell populations reflected in the CellTypist annotation was performed, during which low quality clusters were removed. A single doublet cluster (10087 cells) exhibiting higher Scrublet score as well as co-expression of Macrophage and T cell lineage genes was removed. Since epithelial cells reflected variable, but low level, contaminants in BAL samples, 11282 cells within clusters annotated as epithelial by CellTypist were also excluded before re-clustering leukocytes only. A clustering resolution of 1.4 was applied, before further manual adjustments to either resolve specific cell types annotated by CellTypist, or to resolve observed heterogeneity within neutrophils (0.5 for T and NK cells, 0.3 for neutrophils, 0.15 for conventional DCs and 0.15 to resolve plasmacytoid DCs, mast cells and B cells). The resultant 28 clusters were annotated based on CellTypist analysis against three reference datasets^52-54^ with additional refinements based on marker gene expression (**Supplementary Table 4**) that was essential to identify neutrophils, which were not well represented in these reference datasets.

### Differential abundance

Differential abundance of major cell types across conditions was assessed using the cluster-agnostic statistical tool Milo^55^ via the Python perturbation framework Pertpy v1.0.2^56^, using parameters k=150; p=0.1; and d=50. Milo defines a set of representative neighbourhoods on the *k*-nearest neighbour graph of single cells to model neighbourhood abundance, while accounting for the compositional nature of cell count data. Differentially abundant neighbourhoods were visualized on the full UMAP or mapped back to their majority cell types (those neighbourhoods with >60% cell type labelled) and visualised according to Log_2_ fold change between comparisons.

### Single-cell differential gene expression

Pseudobulk differential gene expression (DGE) analysis was carried out using the decoupler package v 2.1.1^57^ to generate the pseudobulks and the Python implementation of DESeq2, PyDESeq2 v0.5.2, for the differential expression^47,58^. Briefly, gene expression was aggregated per individual and per cell type. Genes were filtered based on minimum expression of n counts in at least 15% of X samples, where X was the smallest group in the comparison and n was defined for each cell type individually. DGE analysis was then computed using Wald tests implemented with PyDESeq2, with models adjusted for sex. Genes were determined to be differentially expressed with a fold change of ≥1.5 and Benjamini–Hochberg-adjusted *P* value of <0.05.

### Inference of cell-to-cell communication from scRNA-seq data

Cell-to-cell interactions were inferred using the R package CellChat v1.1.3, with default parameters. The “population.size” parameter was set to TRUE when computing the inferred interaction between cell subsets. Relative predicted signalling contribution of different ligand–receptor pairs and cell types was quantified as indicated in the relevant figures.

### Gene signature evaluation in scRNA-seq data

Enrichment of gene signatures was scored in cells using the scanpy function sc.tl.score_genes. External gene signatures for evaluation in our scRNA-seq data were obtained from the top 50 genes of a protective T1-T17 T cell signature from NHP granuloma^12^ and from determining common type I IFN response genes from *in vitro* type I IFN-stimulated human neutrophils^59^ and human and mouse macrophages^60^. A random forest model was used to rank genes within the published gene signatures according to their contribution to calculating the enrichment score. A signature of inflammatory macrophages in neutrophil-high TB patients and progressors was obtained by k-means clustering of differentially expressed genes and reduced to the top 50 differentially expressed genes by adjusted *P* value across relevant clusters for use in enrichment analysis.

### Analysis of external scRNA-seq data

Published scRNA-seq data from cynomolgus macaque tuberculosis granulomas^15^ were obtained from the Broad Single Cell Portal (SRA: PRJNA900256) and the processed dataset imported and analysed in Python. Transcriptional signatures derived from our human BAL data were also assessed in our previously published scRNA-seq dataset from lung leukocytes of TB-resistant and -susceptible mouse strains during *M. tuberculosis* infection^19^ (GSE298787) and in bulk RNA-seq data from whole mouse lungs from TB-susceptible mice during *M. tuberculosis* infection with and without IFNAR blockade^19^ (GSE298786).

### Flow cytometry

Approximately 10^6^ live cells per staining panel were washed in PBS, before staining for 30 mins at room temperature with amine-reactive Fixable Blue Viability Dye (Thermo Fisher Scientific; diluted 1 in 500 in PBS). Samples were incubated with 25 mg/ml Human Fc Block (BD Biosciences) diluted in staining buffer (PBS + 2% fetal bovine serum + 2mM EDTA) for 5 minutes at room temperature, before addition – without washing – of cocktails of fluorochrome-conjugated monoclonal antibodies; either for pan-leukocyte analysis or T cell-focused analysis, supplemented with True-Stain Monocyte Blocker (BioLegend) to prevent non-specific binding of tandem fluorochromes to monocytes and macrophages. The pan-leukocyte panel comprised the antibodies (RRID in parentheses): HLA-DR-BV785 (AB_2563461), CD206-APC/Cy7 (AB_2144930), CD45-PERCP/Cy5.5 (AB_893338), CD16-AF700 (AB_2278418), CD11c-PE/Cy7 (AB_389351), CD14-PE (AB_314188), CD15-FITC (AB_314196), CD3-BV510 (AB_2561943), CD56-APC (AB_2563913), CD19-BV650 (AB_2562097) and CD4-PE/Dazzle595 (AB_2565847) from BioLegend and CD8a-BUV737 (AB_2870085) and Siglec 8-BV711 (AB_2872332) from BD Biosciences. The T cell-focused phenotyping panel comprised the antibodies: CCR7-BV711 (AB_2563865), CCR6-PE/Cy7 (AB_10916518), CXCR3-APC (AB_10983064), PD-1-BV650 (AB_2566362), HLA-DR-FITC (AB_314682), CD69-APC/Cy7 (AB_314849), CD45-PERCP/Cy5.5 (AB_893338), CD45RA-PE (AB_314412), CD3-BV510 (AB_2561943), CD56-BV785 (AB_2566059), CD103-PE/Dazzle594 (AB_2716189), CD4-AF700 (AB_571943) or CD4-PE/Cy7 (AB_571959) from BioLegend and CD8a-BUV737 (AB_2870085) from BD Biosciences.

Flow cytometry data were acquired on an X20 analyser (BD Biosciences). Autofluorescence was measured in the 450nm 450/50 channel and used as an analysis parameter. Data were analysed using FlowJo v10 software (BD Biosciences). Populations were defined by hierarchical gating as shown in **Extended Data Figs. 3a and 7a** and expressed as either percentage of total leukocytes (defined as live, CD45^+^, single cells), percentage of a parent population, or as absolute counts (calculated as #cell population/#live single cells x #live cells per ml BAL as counted manually on a haemocytometer.

### Luminex protein assays

BAL supernatants were maintained at -80°C for between 21 and 119 weeks before thawing and sterilising by centrifugation at 10 000 x G through Ultrafree® 0.22 um centrifugal filters (Sigma-Aldrich). Three custom Luminex® Discovery Assays were designed and purchased from Bio-Techne and were performed as per manufacturer’s instructions. Samples were run at a minimum of 1:1 dilution with assay buffer. Data were acquired on a Bio-Plex 200 system (Bio-Rad) and concentration values interpolated from standard curves using Bio-Plex Manager software (Bio-Rad). Inferred values below the bottom standard concentration were ignored and interpreted as below detection limit. Values below the effective limit of detection (bottom standard concentration multiplied by 2, the lowest sample dilution factor of) or identical values close to the effective limit of detection obtained for several samples were interpreted as likely noise and so recorded as below detection limit. Assays consisted of 1: CCL1, CCL2, CCL3*, CCL4, CCL5, CCL13, CCL14, CCL15*, CCL19, CCL20, CCL23*, CCL24, CCL25*, CCL26*, CCL27*, CCL28*, CX_3_CL1*, CXCL2*, CXCL6, CXCL12*; 2: C-reactive protein, CXCL9, G-CSF, GITR*, GM-CSF, Granzyme B, IFN-α, IFN-β*, IFN-γ, IL-1α, IL-1β, IL-RA, IL-2*, IL-3*, IL-4*, IL-5*, IL-10,* IL-11*, IL-12/23p40*, IL-17A*, IL-17C, IL-18, IL-19, IL-21*, IL-28A*, IL-28B*, MMP-1, MMP-12 and TNF-α; 3: CXCL10, CCL18, CXCL16, IL-8, IL-6, MMP-2, MMP-9, MMP-8, CXCL11, CCL17, CCL22. Asterisks in the above lists indicate analytes that were not present above the limit of detection in at least 8 samples and so were not included in the presented analysis. C-reactive protein data were not included in the presented analysis as values were over the limit of detection in several samples. Luminex data were visualised on a row-scaled heatmap generated using the pheatmap R package using clustering method "ward.D2".

### Blinding and randomisation

RNA extraction for bulk RNA-seq was performed in batches in the order of sample collection while blind to final patient group. Single cell RNA-seq runs of pooled samples were performed in parallel to sample collection and so pool composition was determined based on similar sampling date and on similar cellular composition of BAL as determined by flow cytometry, following our preliminary observations that this improved the balance of sequencing coverage across samples within pools. The operator preparing and running scRNA-seq pools was blind to the study group. Bulk and scRNA-seq library preparation was performed through automated core facility pipelines at the Francis Crick Institute by operators blind to the study groups.

### Statistical analysis

Unless otherwise specified, statistical analysis was performed using Prism 10 software (GraphPad) using appropriate statistical tests for the number of groups analysed and the distribution of data, which was determined using Shapiro-Wilk normality tests in Prism 10. Two-tailed tests were used. Descriptive statistics, statistical tests and results are specified in the relevant figures and accompanying legends.

## Data availability

Bulk and scRNA-seq data generated in this study have been deposited in GEO and will be made publicly available upon publication of this work. All code to reproduce the analysis has been deposited at https://github.com/ogarralab/branchett_human_bal_tb_2026 and will be made publicly available with a permanent DOI upon publication of this work.

## Acknowledgements

The authors thank Evangelos Stavropoulos and Alaa Al-Dibouni, of the Immunoregulation and Infection Laboratory at the Francis Crick Institute, for assistance with BAL sample processing; Clare Lloyd and members of the Lloyd Laboratory at Imperial College London, for advice on BAL sample processing and flow cytometry panels; Laure Botella, Izabela Glegola-Madejska and the rest of the Crick Biosafety team for support with work at containment level 3; Phil Hobson, Ana Agua Doce and Kerol Bartolovic of the Crick flow cytometry facility support with flow cytometry at containment level 3; and the Crick Logistics team for fast and secure receipt of BAL samples. The authors additionally thank Gabriella Pountney, nuclear medicine clinical scientist, for support with providing pseudonymised PET-CT images; Kumaran Balasundaram, clinical trials physician, for support with chest X-ray scoring; and Maes Mailis, Sharon Cox and Susanne Howes from the UK-Health Security Agency TB Unit for their support providing data of lineage and relatedness from whole genome sequencing of *M. tuberculosis* culture positive specimens in the study cohort.

This work was funded by a Wellcome Investigator Award to AOG (WT 215628/Z/19/Z) which financially supported all consumables and salaries of WJB, JWK, JL, IN and JS, although WJB, JS, PC, HS, KAW, RJW and AOG are employed by the Francis Crick Institute, which receives core funding from Cancer Research UK, the UK Medical Research Council and Wellcome. AOG and lab, are supported by the Francis Crick Institute (CC2084). This study has been delivered through the National Institute for Health and Care Research (NIHR) Leicester Biomedical Research Centre (NIHR203327). The views expressed are those of the authors and not necessarily those of the funder, the NIHR or the Department of Health and Social Care.

## Author contributions

AOG and PH designed the study with input from RJW; JWK managed participant recruitment and follow-up, with assistance from JL and IN; JWK, RV and PH performed bronchoscopy to obtain BAL and performed the PET-CT scans; JWK performed analysis of PET-CT images, under the supervision of AK; WJB processed and analysed all BAL samples and coordinated data with clinical parameters; JS performed bulk and scRNA-Seq analysis, under the guidance of AOG and WJB; PC performed initial scRNA-Seq analysis and accessed and analysed published non-human primate scRNA-Seq data; KAW helped design Luminex experiments, performed Luminex assays and helped with interpretation of data; WJB and JS prepared data visualisations and figures under AOG supervision; WJB, AOG, JWK and PH designed the manuscript structure including figures and wrote the manuscript, with substantial input from RJW, input from KAW and specific input from JS, PC and HS.

## Competing interests

The authors declare no competing interests.

## Materials & Correspondence

Requests for materials and correspondence should be addressed to Dr Anne O’Garra, Principal Group Leader, the Francis Crick Institute.

## Extended Data Figures

**Extended Data Fig. 1.**
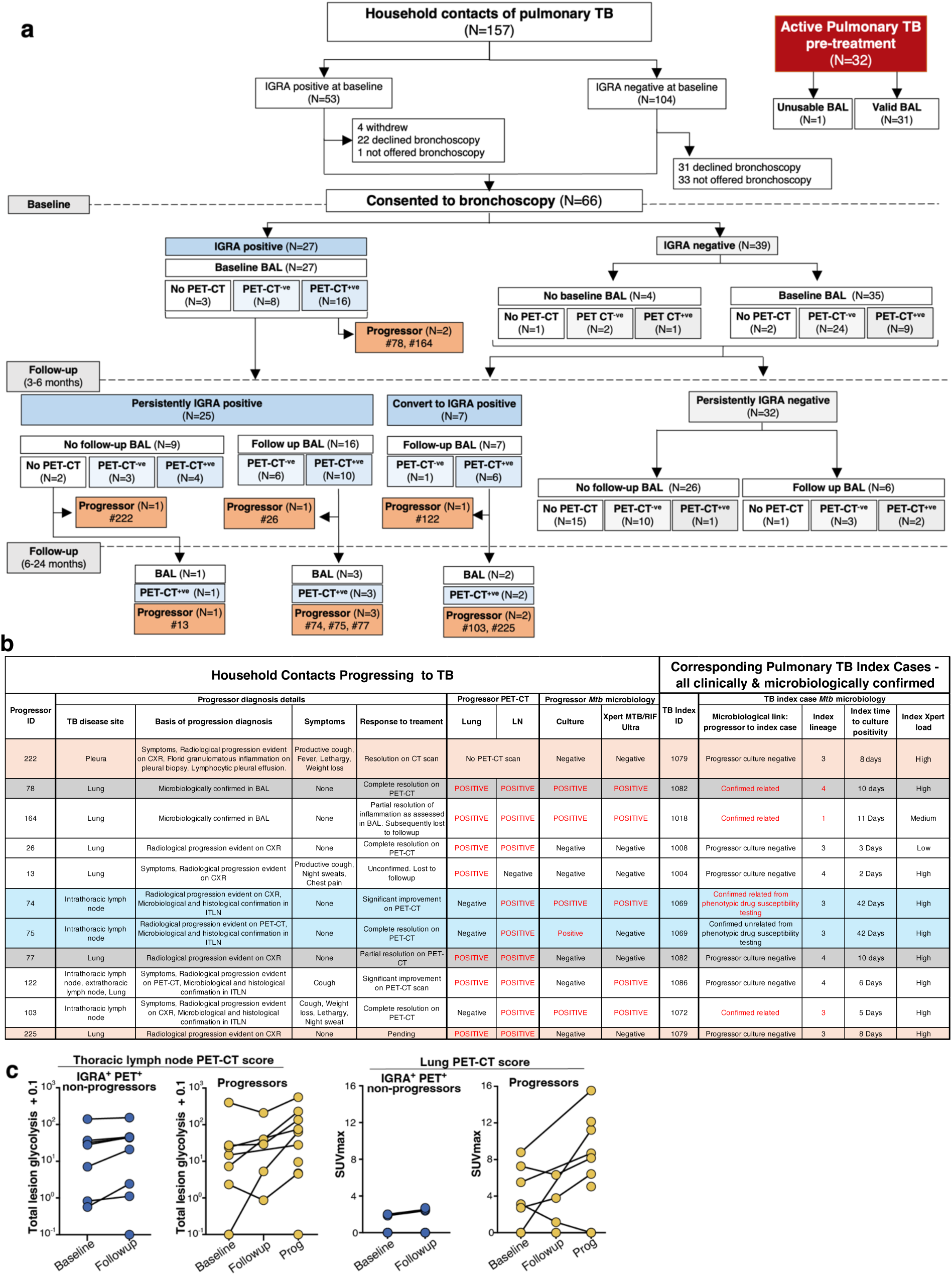
Study overview and details on progressors. **a.** Expanded flow chart showing recruitment, sampling and TB progression events in the study. **b.** Details on contacts who progressed to TB and respective TB index cases. Coloured rows indicate progressors from the same index case. ITLN, intrathoracic lymph node. **c**. PET-CT scores from thoracic lymph node and lung in progressors and IGRA^+^ non-progressors presenting with positive thoracic PET-CT. Total lesion glycolysis (TLG) represents combined metabolic activity from all positive lymph nodes (SUV_mean_ x metabolic volume). SUV_max_ was determined from the highest avid lung lesion. Data are shown from baseline and follow-up visits and at progression, where relevant, with lines connecting individuals. See also **Supplementary Table 1** and **Supplementary Fig. 1**.

**Extended Data Fig. 2.**
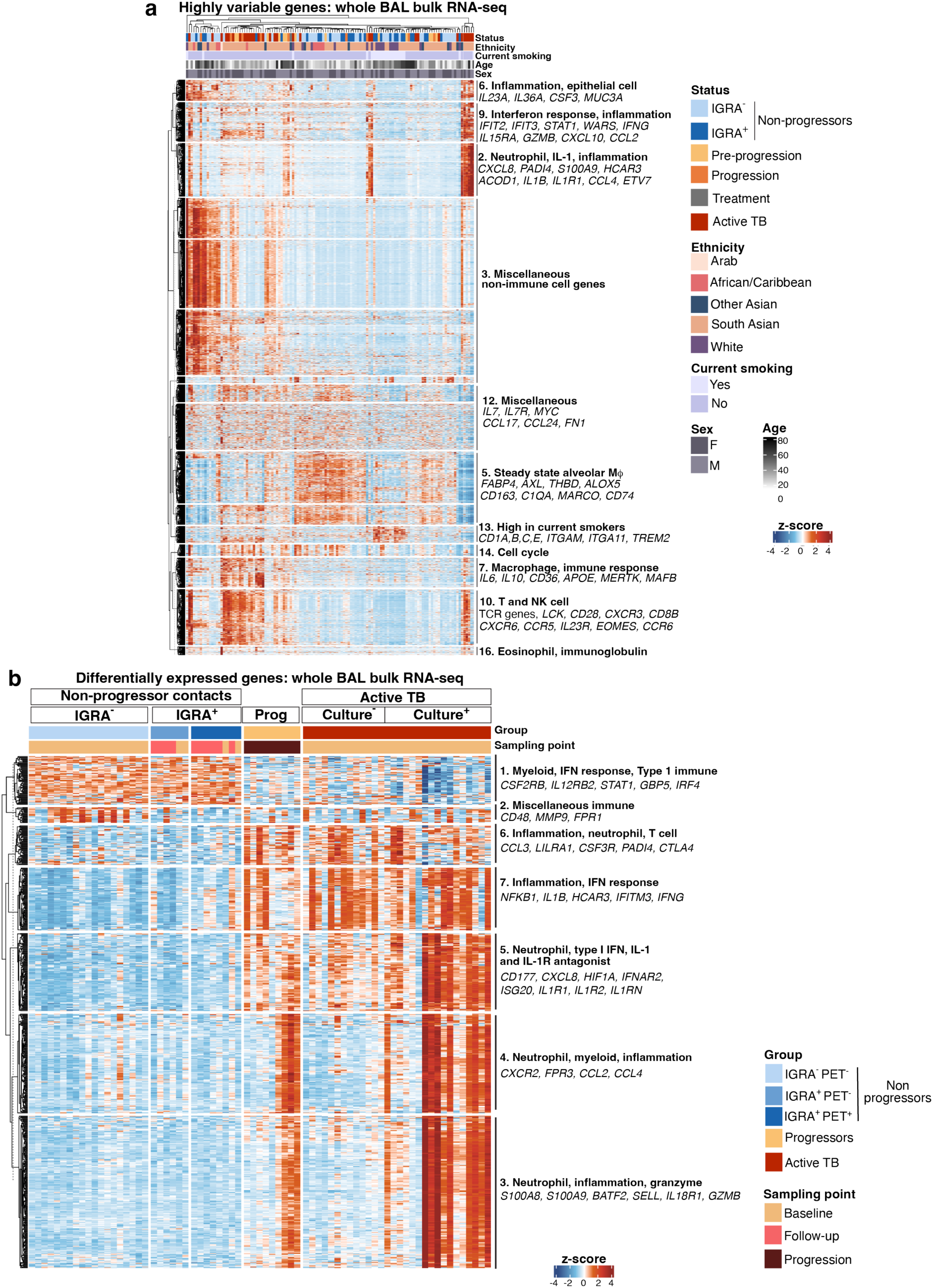
Global transcriptional changes in airway cells of TB patients and household contacts by bulk RNA-seq. Unsupervised Ward’s clustering of highly variable genes among all BAL samples for which bulk RNA-seq was obtained, before excluding any outliers and including multiple timepoints for individuals, where available (**Supplementary Table 1a-b**). **b.** k-means clustering of total differentially expressed genes combined from comparisons between i) IGRA^+^ PET-CT^-^ (N=6) and IGRA^-^PET-CT^-^ (N=19) non-progressors; ii) IGRA^+^ PET-CT^+^ (N=8) and IGRA^-^PET-CT^-^ non-progressors and iii) IGRA^+^ PET-CT^+^ non-progressors and progressors at progression (N=9); clustered across samples from all groups, including BAL culture-positive (N=13) and culture-negative (N=17) active TB, after excluding outlier samples as detailed in **Methods** and **Supplementary Tables 1a-b**. In both panels, cluster numbers are stated and representative genes and associated pathways and cell types from each cluster are listed (see also **Supplementary Tables 2** and **3**).

**Extended Data Fig. 3.**
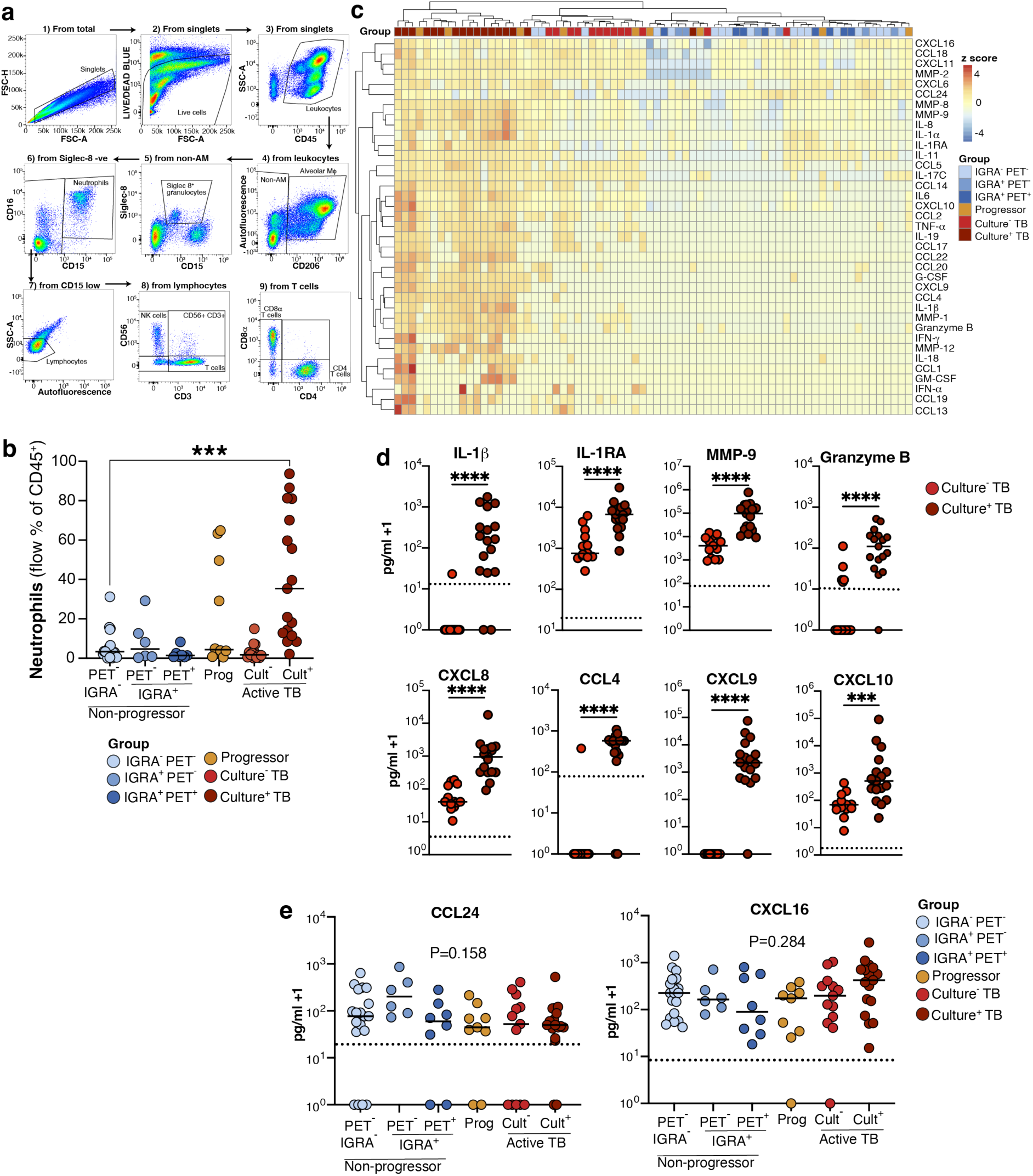
Bronchoalveolar lavage flow cytometry and protein multiplex assays. **a.** Flow cytometry gating strategy for the pan-immune cell panel used to quantify neutrophils (CD15^hi^ CD16^+^ CD206^low-neg^). **b.** Percentage of neutrophils within total live CD45^+^ BAL leukocytes as determined by flow cytometry. Points show data from individual participants with lines at the median; IGRA^-^ PET-CT^-^ (N=19), IGRA^+^ PET-CT^-^ (N=6), IGRA^+^ PET-CT^+^ (N=7), Progressors at progression (N=10); BAL culture^-^ TB (N=13) BAL culture^+^ TB (N=17). Statistics: Kruskal-Wallis with Dunn’s post hoc test; ***, *P* < 0.001. **c.** Row-scaled heat-map showing Ward’s clustering of samples based on analyte concentrations in BAL supernatants as determined by Luminex multiplex immunoassays. **d-e.** Concentrations of selected protein analytes in BAL supernatants, with dashed lines indicating the effective limit of detection. Values of zero indicate no detection above this limit. Points show data from individual participants with lines at the median. IGRA^-^ PET-CT^-^ (N=19), IGRA^+^ PET-CT^-^ (N=6), IGRA^+^ PET-CT^+^ (N=8), Progressors at progression (N=9) and BAL culture-negative (N=13) and - positive (N=17) TB patients. Statistics: **d.** Mann-Whitney test; ***, *P*<0.001; ****, *P*<0.0001 **e.** Kruskal-Wallis test.

**Extended Data Fig. 4.**
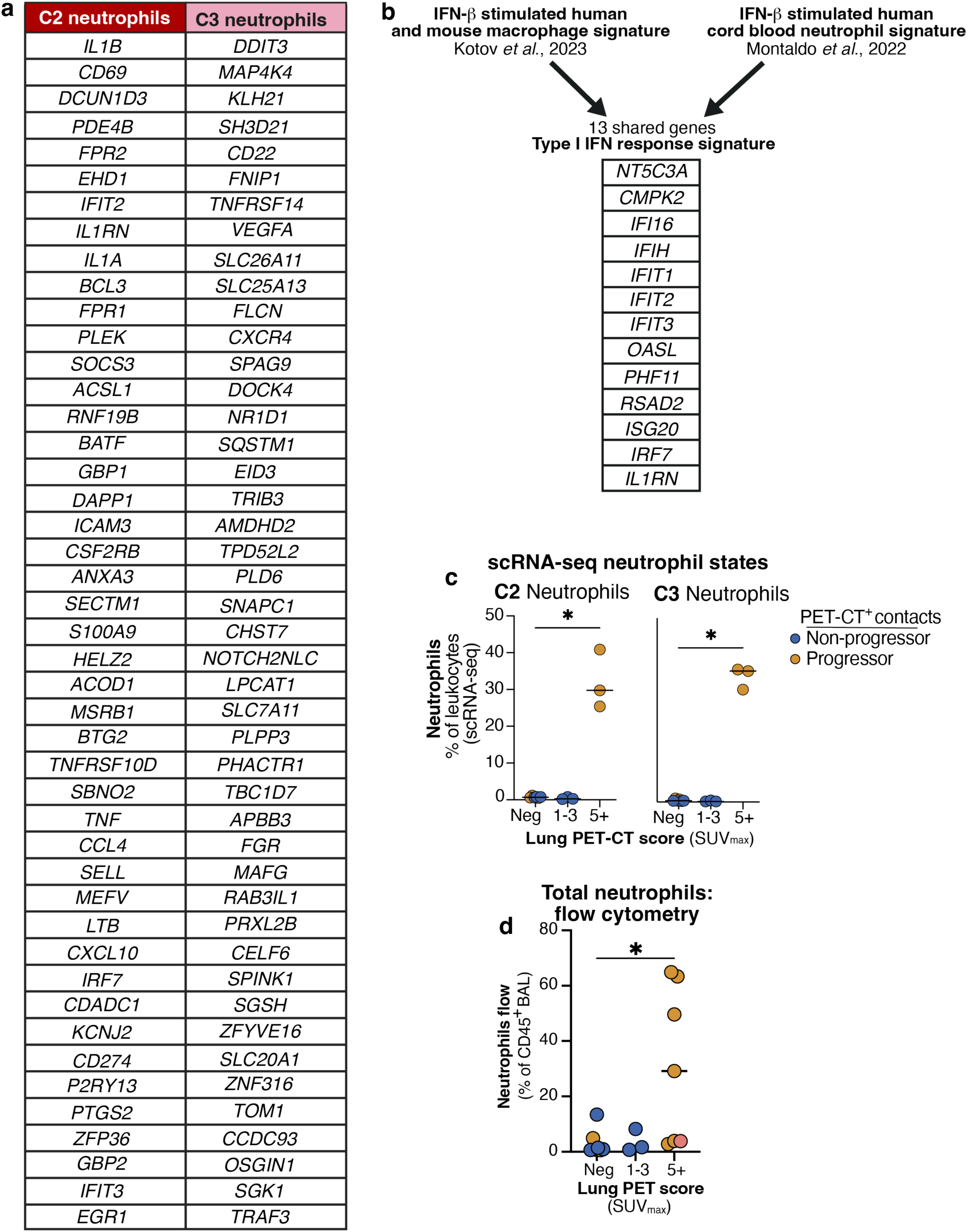
Analysis of airway neutrophils by scRNA-seq and flow cytometry. **a.** Lists of top 50 differentially expressed genes between cluster 2 (C2) and C3 neutrophils, ranked by adjusted *P* value. **b.** Schematic showing selection of a 13 gene myeloid type I IFN response signature by combination of signatures from the indicated publications. **c.** Frequency of C2 and C3 neutrophils as a percentage of total BAL leukocytes by scRNA-seq in all PET-CT^+^ TB contacts, binned by lung PET-CT status and coloured by progressor status (N=3 to 6 per lung PET-CT bin). **d.** Percentage of total neutrophils by flow cytometry as a percentage of total live CD45^+^ BAL leukocytes as determined by flow cytometry, plotted as for **c** (N=3 to 7 per lung PET-CT bin). Points in **c-d** represent individual contacts with lines at the median. Statistics: Kruskal-Wallis test with Dunn’s post-hoc test; *, *P* <0.05.

**Extended Data Fig. 5.**
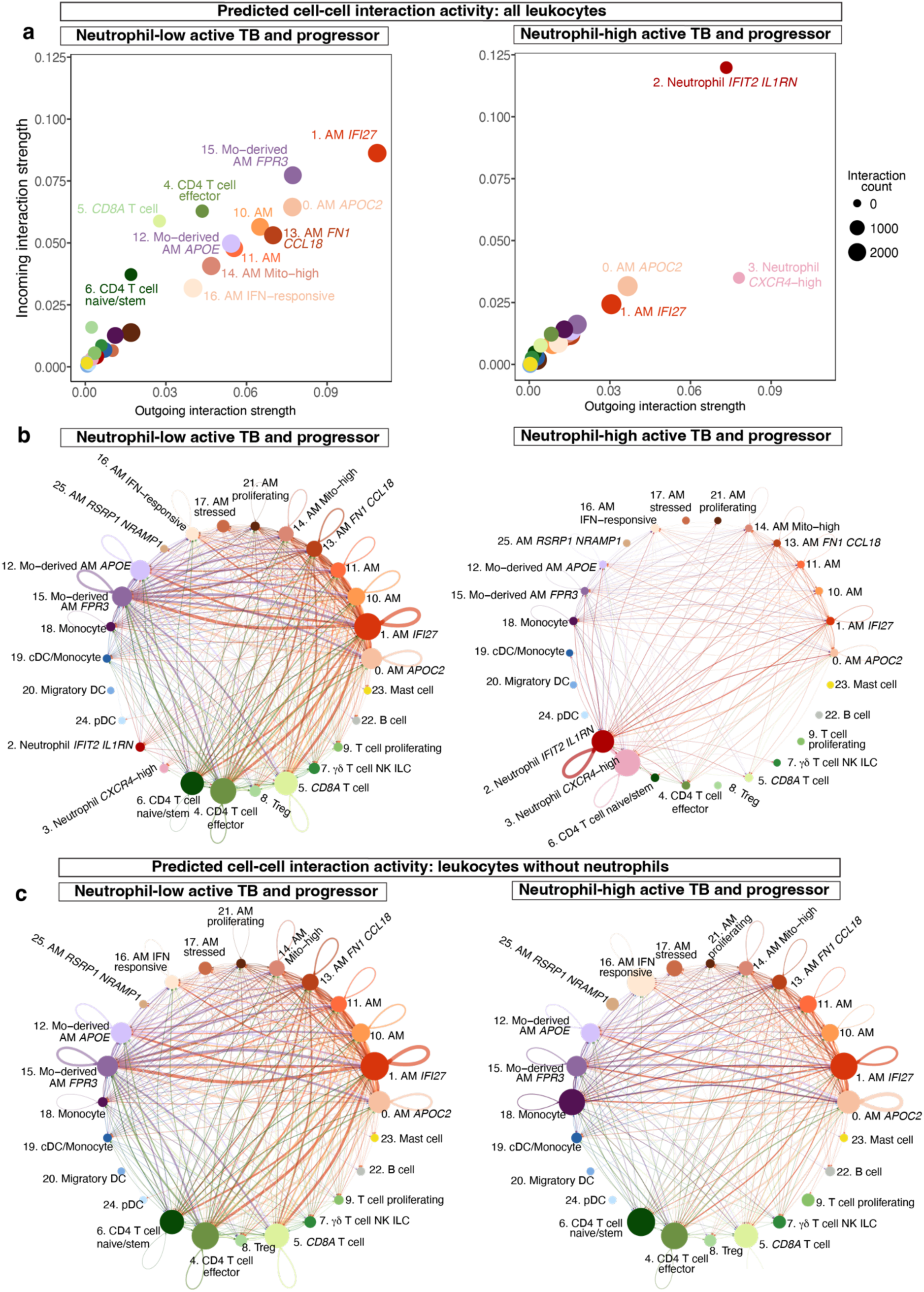
Predicted immune interactions in airways of neutrophil-low and –high TB patients and progressors. **a-c.** CellChat inferred receptor-ligand interactions analysis comparing Neutrophil-high to Neutrophil-low TB patients and progressors, defined, as having ≥ or < 15% neutrophils, respectively, as a percentage of live, CD45^+^ BAL leukocytes by flow cytometry. **a.** Outgoing and incoming signal strength per scRNA-seq cluster. **b-c.** Predicted total interaction strength between scRNA-seq clusters in each condition, shown either with all cell types included (**b**) or with neutrophils excluded from the visualization (**c**). Circle sizes are proportional to the abundance of each cluster, lines are coloured according to the signal-sending population and line thickness represents the relative predicted strength of interactions. Data are averaged from a total of 16 neutrophil-low and 11 neutrophil-high samples.

**Extended Data Fig. 6.**
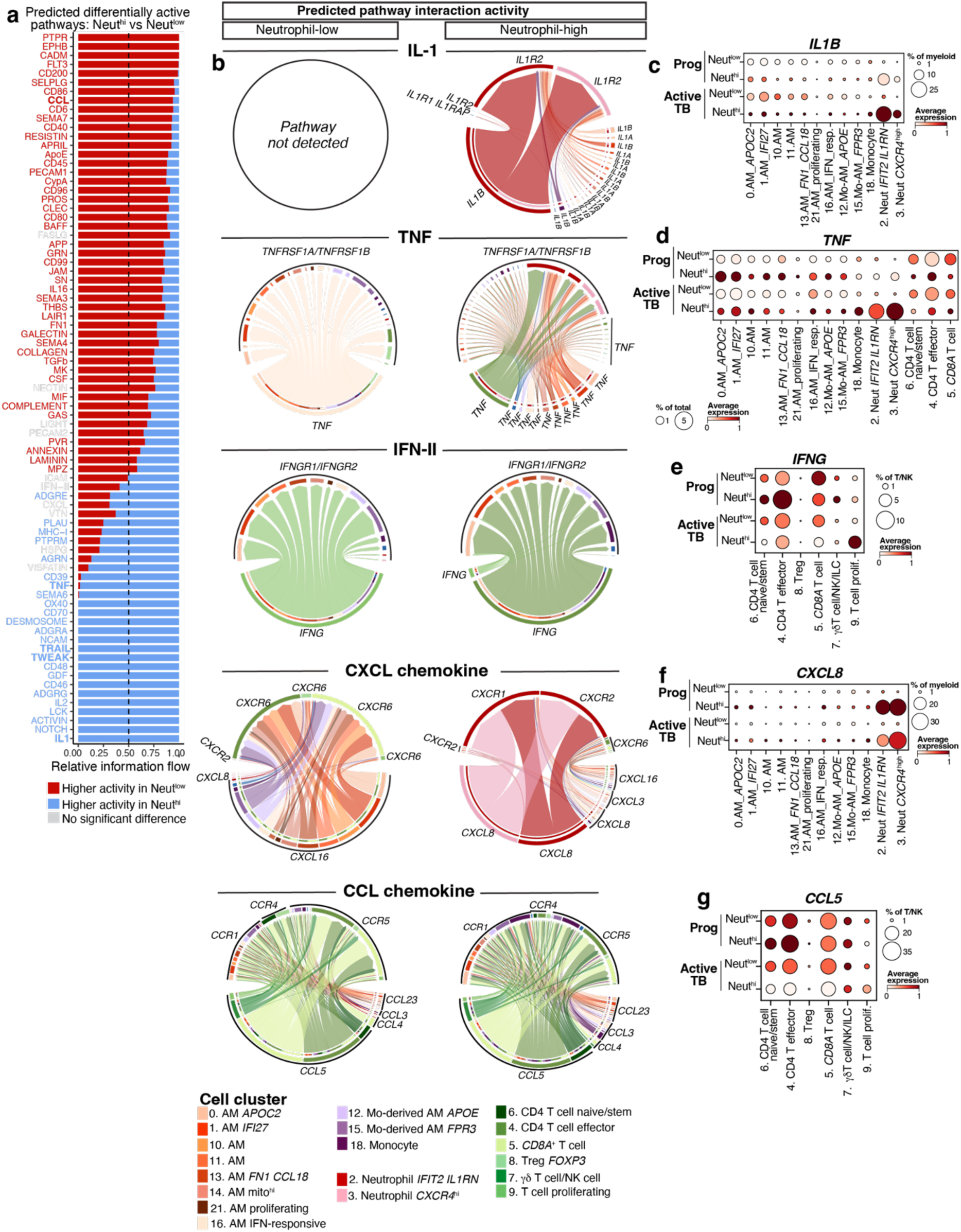
Predicted differentially active pathways in neutrophil-low and – high TB patients and progressors. **a.** Bar plot showing predicted differentially active pathways by CellChat in neutrophil-low (N=16) and neutrophil-high (N=11) TB patients and progressors. The neutrophil low and -high states were defined, as having < or ≥15% neutrophils, respectively, as a percentage of live, CD45^+^ BAL leukocytes by flow cytometry. Significant differences in predicted pathway activity were determined by permutation analysis in CellChat, with *P* <0.05 taken as significant. **b.** Plots showing the predicted signalling between the indicated scRNA-Seq clusters and receptor-ligand pairs in each condition. Data shown are averaged from a total of 16 neutrophil-low and 11 neutrophil-high samples. **c-g.** Dot plots showing expression of key ligand genes across relevant scRNA-seq clusters in neutrophil-low progressors (N=3), neutrophil-high progressors (N=3); neutrophil-low TB patients (N=13) and neutrophil-high TB patients (N=8).

**Extended Data Fig. 7.**
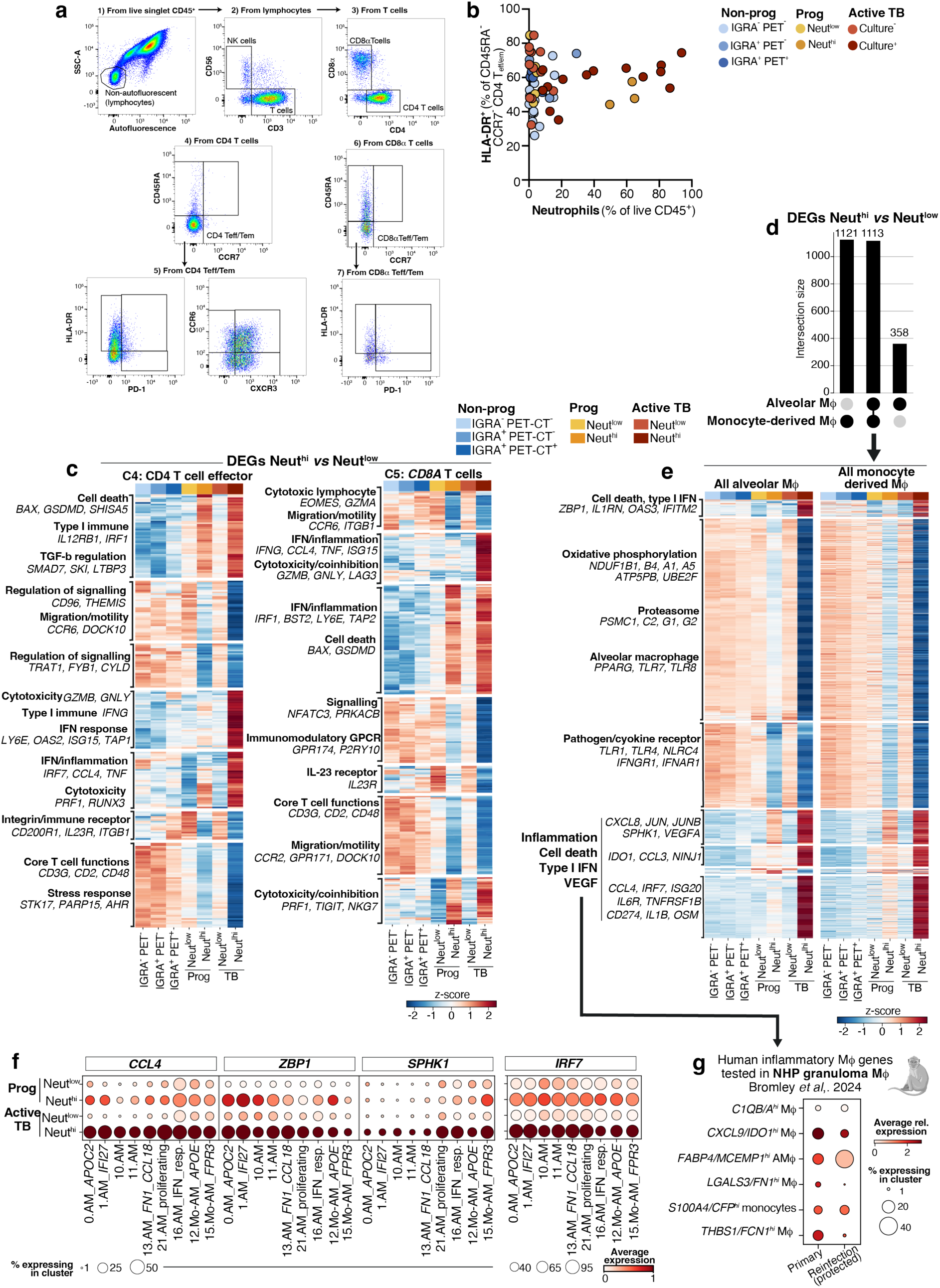
Airway immune signatures accompanying the neutrophil-high state. **a.** Flow cytometry gating strategy for the T cell focused panel, used to identify different T cell phenotypic and activation markers. **b.** HLA-DR^+^ CD4^+^ CD45RA^-^ CCR7^-^ T_eff/em_ as a percentage of total CD4^+^ CD45RA^-^ CCR7^-^ T_eff/em_, plotted against neutrophil frequency in BAL by flow cytometry. Data are shown from IGRA^-^ PET-CT^-^ (N=15), IGRA^+^ PET-CT^-^ (N=6), IGRA^+^ PET-CT^+^ (N=7), neutrophil-low progressors (N=6), neutrophil-high progressors (N=3), BAL culture-negative TB patients (N=12) and BAL culture-positive TB patients (N=14). **c.** Heatmaps showing k-means clustering of differentially expressed genes from T cell populations in scRNA-seq data from neutrophil-low and -high TB patients and progressors defined as ≥ or < 15% neutrophils as a percentage of live, CD45^+^ BAL leukocytes by flow cytometry, respectively. **d.** Upset plot showing shared and unique differentially expressed gene numbers between alveolar macrophage and monocyte-derived macrophage analyses comparing Neutrophil-high and -low TB patients and progressors. **e.** Heatmap showing k-means clustering of shared differentially expressed genes from **d**. **f.** Expression of additional representative genes increased in airway macrophage from neutrophil-high compared to neutrophil-low active TB patients and progressors. Data shown in **c-f** are averaged from all cells from all samples per group: neutrophil-low progressors (N=3); neutrophil-high progressors (N=3) and neutrophil-low (N=13) and neutrophil-high (N=8) TB patients. **g.** Expression of a 50 gene signature of human airway inflammatory macrophages shown in macrophage clusters from cells averaged from NHP granulomas of primary *M. tuberculosis* infection (N=10) compared to reinfection (N=8).

**Extended Data Fig. 8.**
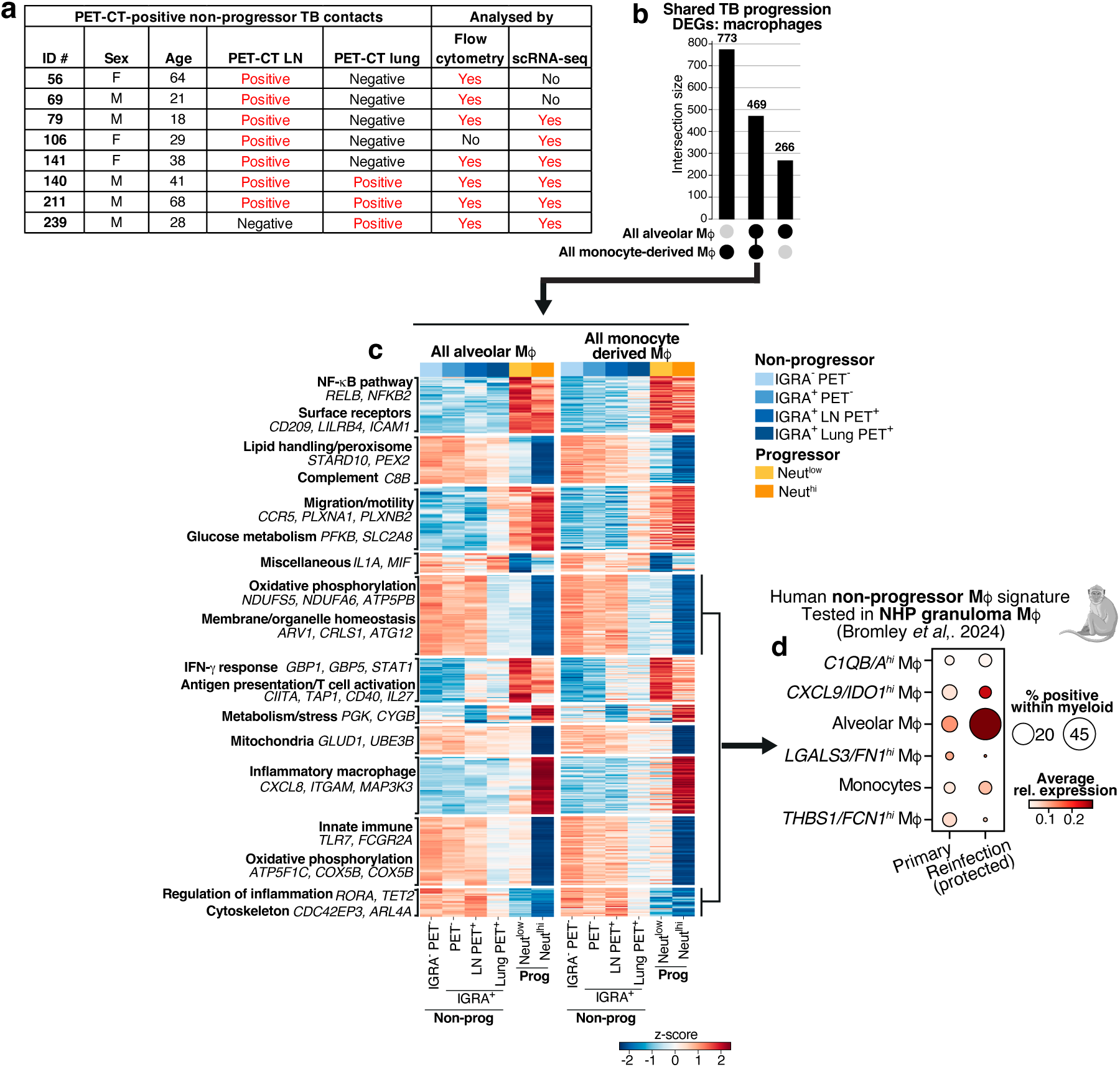
Reduced expression of steady state macrophage transcriptional signatures with progression to TB. **a.** Table showing details of intrathoracic LN and lung parenchyma PET-CT status in PET-CT^+^ IGRA^+^ non-progressors. LN PET-CT^+^ IGRA^+^ non-progressors were defined as those with positive LN PET-CT signal, without positive lung PET-CT signal. **b.** Upset plot showing shared and distinct differentially expressed genes in alveolar and monocyte-derived macrophage scRNA-seq clusters combined from comparisons between progressor and non-progressor sub-groups (detailed in **Supplementary Table 7**). **c.** Heatmap showing k-means clustering of shared alveolar and monocyte-derived macrophage differentially expressed genes across IGRA^-^ PET-CT^-^ (N=9), IGRA^+^ PET-CT^-^ (N=5), IGRA^+^ LN PET-CT^+^ (N=3), IGRA^+^ lung PET-CT^+^ (N=3), neutrophil-low progressors (N=3), and neutrophil-high progressors (N=3). **d.** Expression of a 50 gene signature of human airway macrophages that decreased in TB progressors, shown in macrophage scRNA-seq clusters from NHP granulomas in primary (N=10) *M. tuberculosis* infection compared to reinfection (N=8).

**Extended Data Fig. 9.**
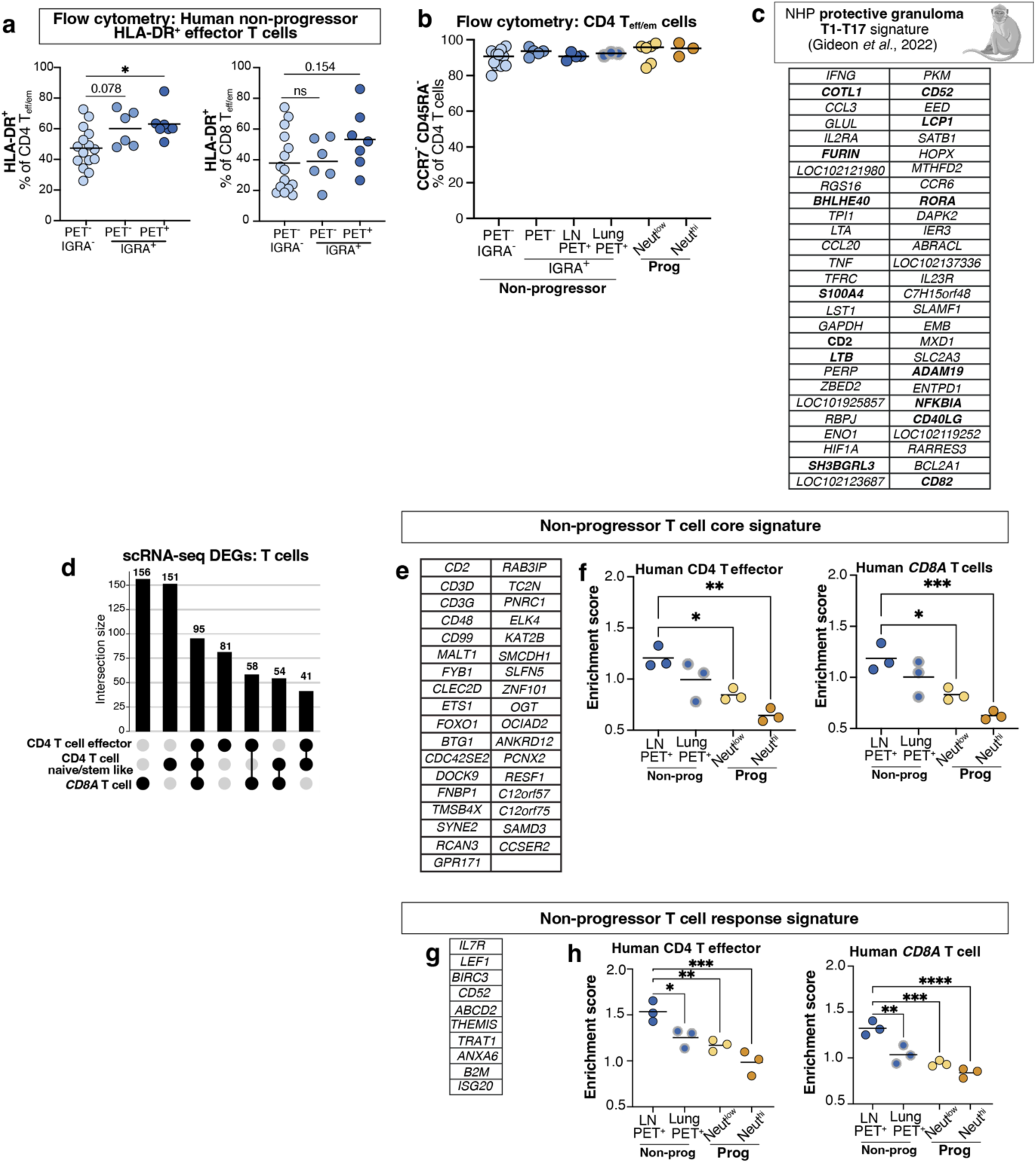
T cell signatures of non-progression in human TB contacts. **a-b.** T cell marker expression as determined by flow cytometry in IGRA^-^ PET-CT^-^ (N=15), IGRA^+^ PET-CT^-^ (N=6), IGRA^+^ PET-CT^+^ (N=7), neutrophil-low progressor (N=5) and neutrophil-high (N=3) progressor BAL. **a.** Percentage of T_eff/em_ CD4 and CD8 T cells expressing HLA-DR. Points show data from individual contacts with lines at the median. Statistics: Kruskal-Wallis test with Dunn’s *post hoc* test; *, *P*<0.05. **b.** Percentage of total CD4 T cells with a CD45RA^-^CCR7^-^ T_eff/em_ surface phenotype. Points show data from individual participants with lines at the mean. **c.** List of genes derived from the published protective NHP signature as used in **Fig. 5c**. Genes in bold are those contributing ≥1% to the observed enrichment in our human BAL CD4 T effector cells. **d.** Upset plot showing shared and distinct differentially expressed genes in the major T cell scRNA-seq clusters combined from comparisons between progressor and non-progressor sub-groups (detailed in **Supplementary Table 7**); IGRA^-^ PET-CT^-^ (N=9), IGRA^+^ PET-CT^-^ (N=5), IGRA^+^ LN PET-CT^+^ (N=3), IGRA^+^ lung PET-CT^+^ (N=3), neutrophil-low progressors (N=3) and neutrophil-high progressors (N=3). **e,g.** Lists of genes within the indicated T cell signatures as shown in **Fig. 5**. **f,h.** Mean enrichment scores of the indicated signatures across all CD4 effector and CD8A T cells in individual contacts. Points show data from individual participants with lines at the mean (N=3 per group). Statistics: one-way ANOVA with Sidak’s post-hoc test; *, *P*<0.05; **, *P*<0.01; ***, *P*<0.001; ****, *P*<0.0001.

**Extended Data. Fig. 10.**
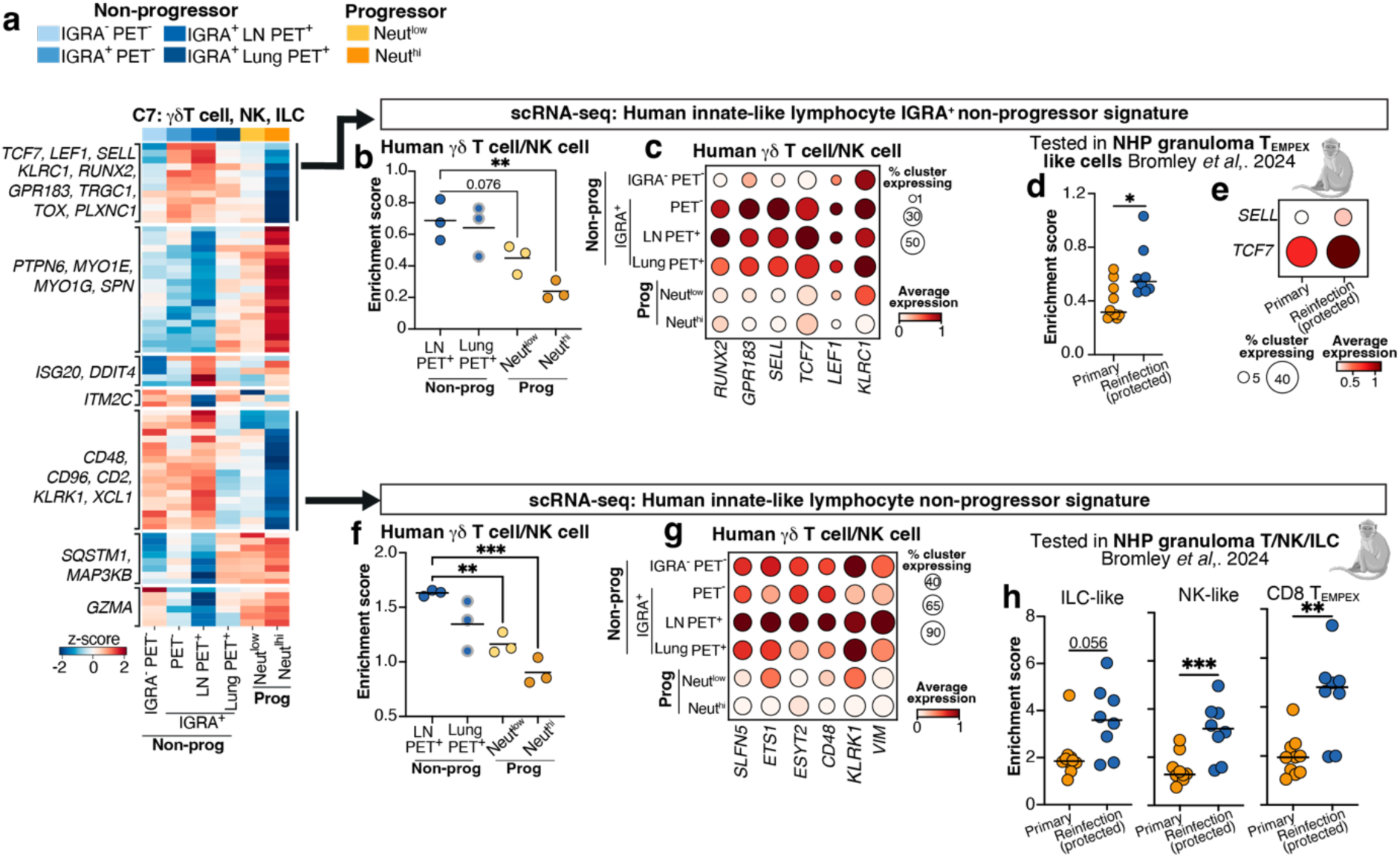
Innate-like lymphocyte transcriptional signatures of non-progression from human TB contacts. **a.** Heatmap showing k-means clustering of differentially expressed genes from the γδT cell, NK, ILC scRNA-seq cluster, combined from comparisons between progressor and non-progressor sub-groups (detailed in **Supplementary Table 7**); IGRA^-^ PET-CT^-^ (N=9), IGRA^+^ PET-CT^-^ (N=5), IGRA^+^ LN PET-CT^+^ (N=3), IGRA^+^ lung PET-CT^+^ (N=3), neutrophil-low progressors (N=3) and neutrophil-high progressors (N=3). **b,f.** Mean enrichment scores of the indicated signatures across all γδT cell, NK, ILC cells in the indicated groups. Points show data from individual contacts with lines at the mean (N=3 per group). Statistics: one-way ANOVA with Sidak’s post-hoc test; **, *P*<0.01; ***, *P*<0.001. **c,g.** Expression of representative genes of the indicated signatures shown from all γ8T cell, NK, ILC averaged across all samples per group. **d,h.** Mean enrichment scores of human non-progressor γδT cell, NK, ILC signatures shown in the indicated scRNA-seq clusters from NHP granulomas in primary (N=10) *M. tuberculosis* infection compared to reinfection (N=8). Points show individual granulomas with lines at the median; statistics: Mann-Whitney test; *, *P*<0.05; **, *P*<0.01; *P*<0.001. **e.** Expression of stem-like T cell genes shown in the CD8 T_em/pex_-like cluster from NHP granulomas in primary (N=10) *M. tuberculosis* infection compared to reinfection (N=8).

## Supplementary Information

**Supplementary Fig. 1. PET-CT scan images from all contacts who received bronchoscopy**

a. All household contacts progressing to TB

b. All IGRA^+^ non-progressor contacts

c. All IGRA^-^ non-progressor contacts

**Supplementary Table 1. Details of study participants and samples obtained**

a. All TB household contacts, including progressors

b. All TB patients

c. Full details on progressing contacts

**Supplementary Table 2. Lists of genes in the hierarchically clustered heat-map of whole BAL bulk RNA-seq data in Extended Data 2a**

**Supplementary Table 3. Lists of genes in the k-means clusters of whole BAL bulk RNA-seq differential expression analysis in Extended Data 2b**

**Supplementary Table 4. Details of scRNA-seq cluster annotations**

**Supplementary Table 5. Numbers and proportions of scRNA-seq clusters per sample**

**Supplementary Table 6. Lists of differentially expressed genes in scRNA-seq BAL leukocyte populations from neutrophil-high compared to neutrophil-low comparisons TB patients and progressors** (accompanies Fig. 4)

**Supplementary Table 7. Lists of differentially expressed genes in scRNA-seq BAL leukocyte populations from comparisons between sub-groups of non-progressor and progressor contacts** (accompanies Fig. 5)

## Notes

### Competing Interest Statement

The authors have declared no competing interest.

