## Supplementary Fig. 1 for "Airway immune signatures of protection and disease progression in recent human tuberculosis household contacts"

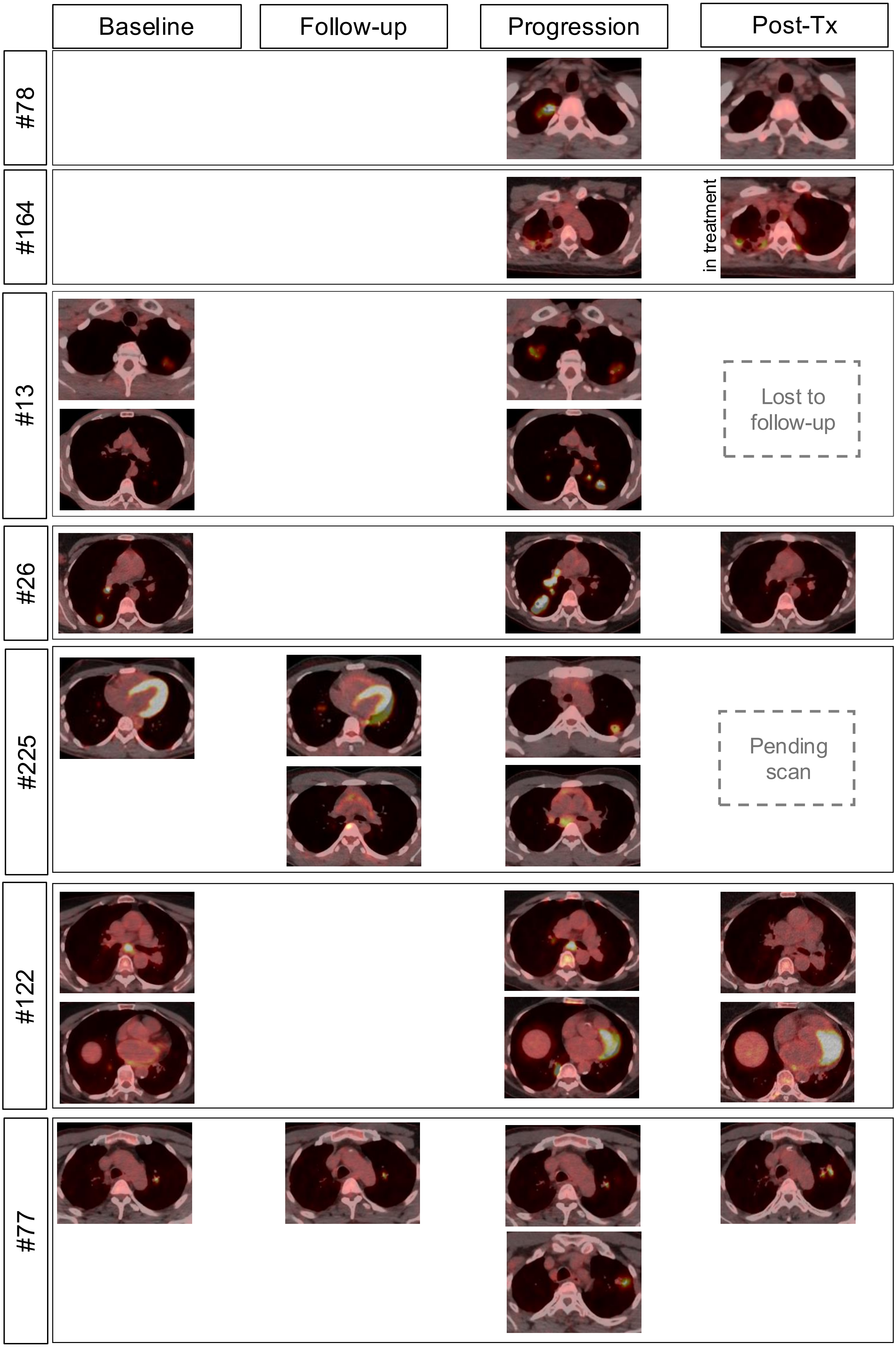


**Supplementary Information Fig. 1a.** Thoracic PET-CT scans of progressing contacts

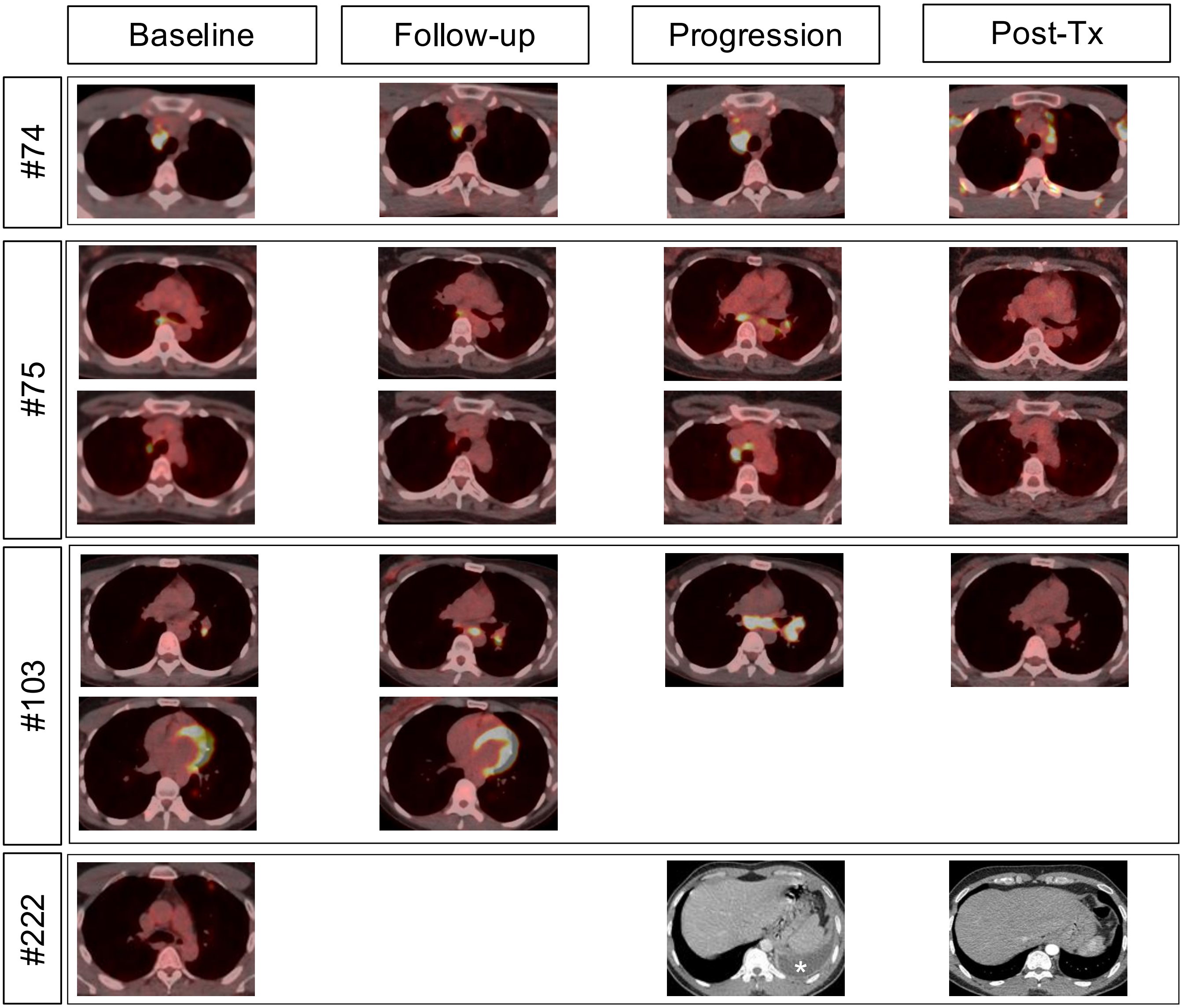


**Supplementary Information Fig. 1a (continued).** Thoracic PET-CT or CT only images from progressing contacts. *, indicates pleural effusion on CT of one progressor that resolved with treatment.

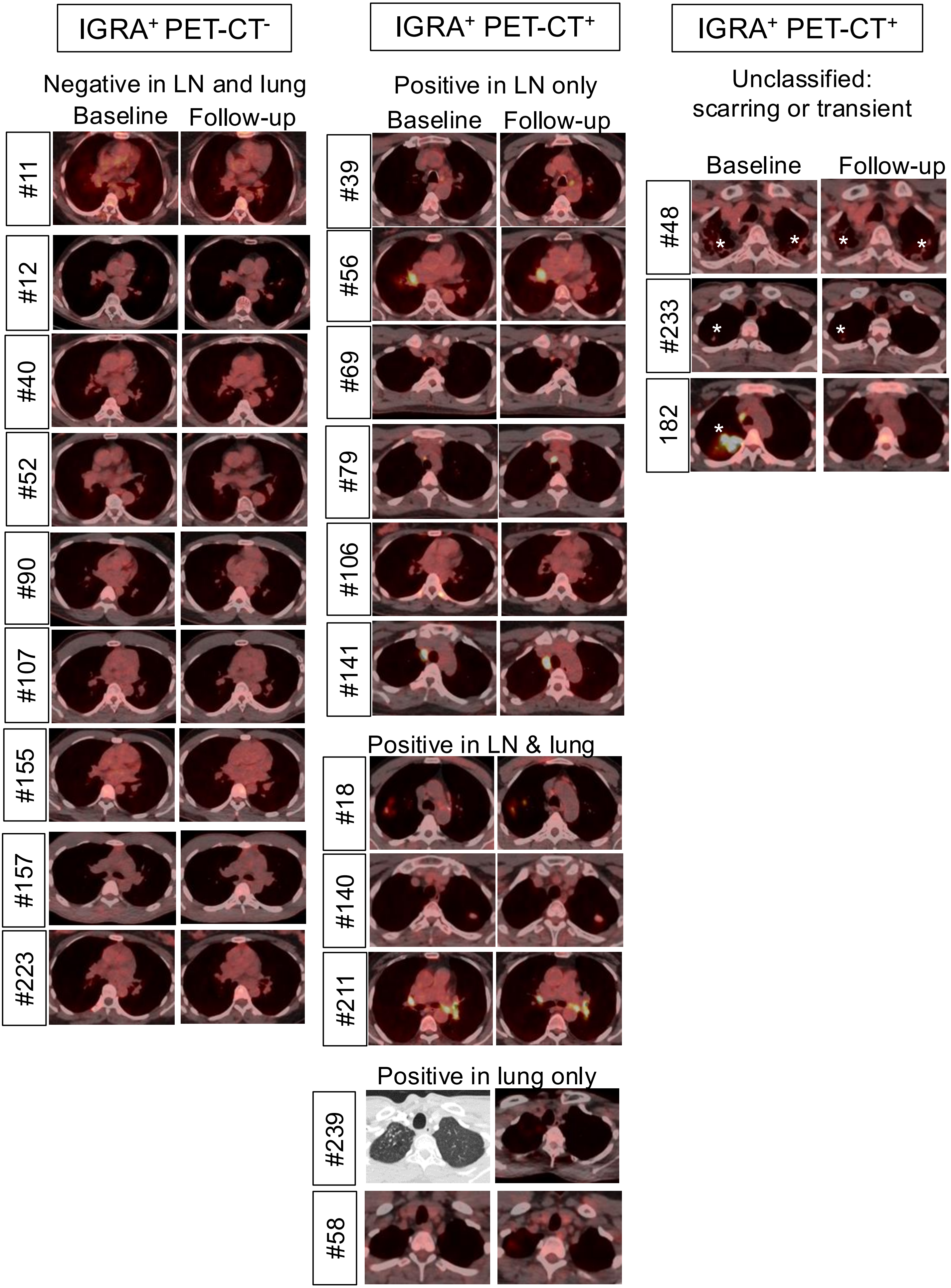


**Supplementary Information Fig. 1b.** Thoracic PET-CT or CT only images in IGRA+ non-progressors. *, indicates scarring or transient positive PET-CT signal

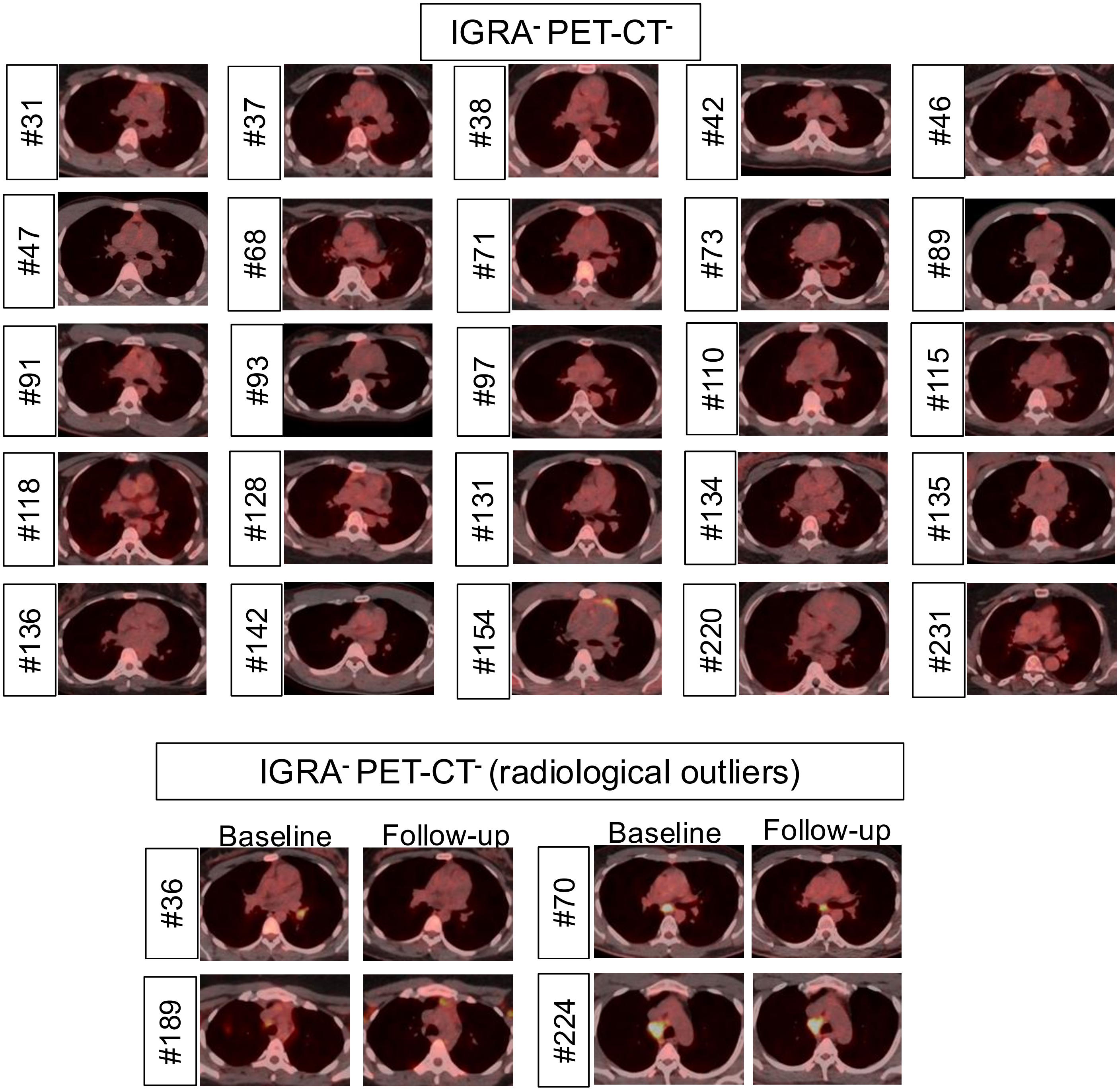


**Supplementary Information Fig. 1c** Thoracic PET-CT images in IGRA$^-$ non-progressors.
